# C-C bond cleavage and generation in TCA cycle driven by radicals and peroxo-bridge like interaction

**DOI:** 10.1101/2024.11.06.622199

**Authors:** Lin Yang, Yijun Fang, Weimin Gong

## Abstract

Tricarboxylic acid (TCA) cycle is one of the most important and common metabolic pathways in different kingdoms of life, in which citrate synthase catalyzes the first irreversible reaction and the rate-limiting step. Reversed oxidative tricarboxylic acid (roTCA) cycle is a newly discovered autotrophic CO_2_ fixation pathway based on the reversal of TCA cycle, in which citrate synthase catalyzes the thermodynamically unfavorable citrate cleavage reaction. In this work, we proposed that radicals promote C-C bond cleavage of citrate in roTCA cycle based on series intermediate catalytic state of citrate synthases from *Desulfurella acetivoran* (*Da*CS) and *Thermosulfidibacter takaii* (*Tt*CS) obtained by electron paramagnetic resonance (EPR) and X-ray crystallography. We also proposed that the peroxo-bridge like interaction promotes C-C bond generation between acetyl group and oxaloacetate in citrate synthesis reaction based on intermediate state of human citrate synthase by time-resolved characteristic Raman spectroscopy and X-ray diffraction methods. Results also indicate these mechanisms would be widespread among conserved citrate synthases.

## Main

CO_2_ fixation, one of the most important biosynthetic processes in nature^1^, is the basic source of organic carbon in the biosphere. Autotrophs convert light and thermal energy into chemical energy though CO_2_ fixation and provide it to all other organisms within ecosystem^2–3^. The reducing tricarboxylic acid (rTCA) cycle is one of the earliest autotrophic CO_2_ fixation processes known^1,3^. As reverse operation of the oxidative TCA (oTCA) cycle, rTCA cycle requires several characteristic enzymes to modify the three irreversible reactions in oTCA cycle under physiological conditions (Supplementary Fig. 1b)^1–2^. Reversed oxidative tricarboxylic acid (roTCA) cycle, a variant of rTCA cycle identified in two thermophilic anaerobic bacteria (*Desulfurella acetivorans*^4^ and *Thermosulfidibacter takaii*^5^), uses citrate synthase in oTCA cycle to catalyze the reversal of citrate synthesis reaction which was considered to be irreversible in oTCA cycle^4–6^ (Fig. 1a). Citrate synthase in TCA cycle, catalyzes the only C-C bond generation reaction^7–9^ without metal cofactor^9–11^, has aroused widespread interests about its mechanism.

**Fig. 1:**
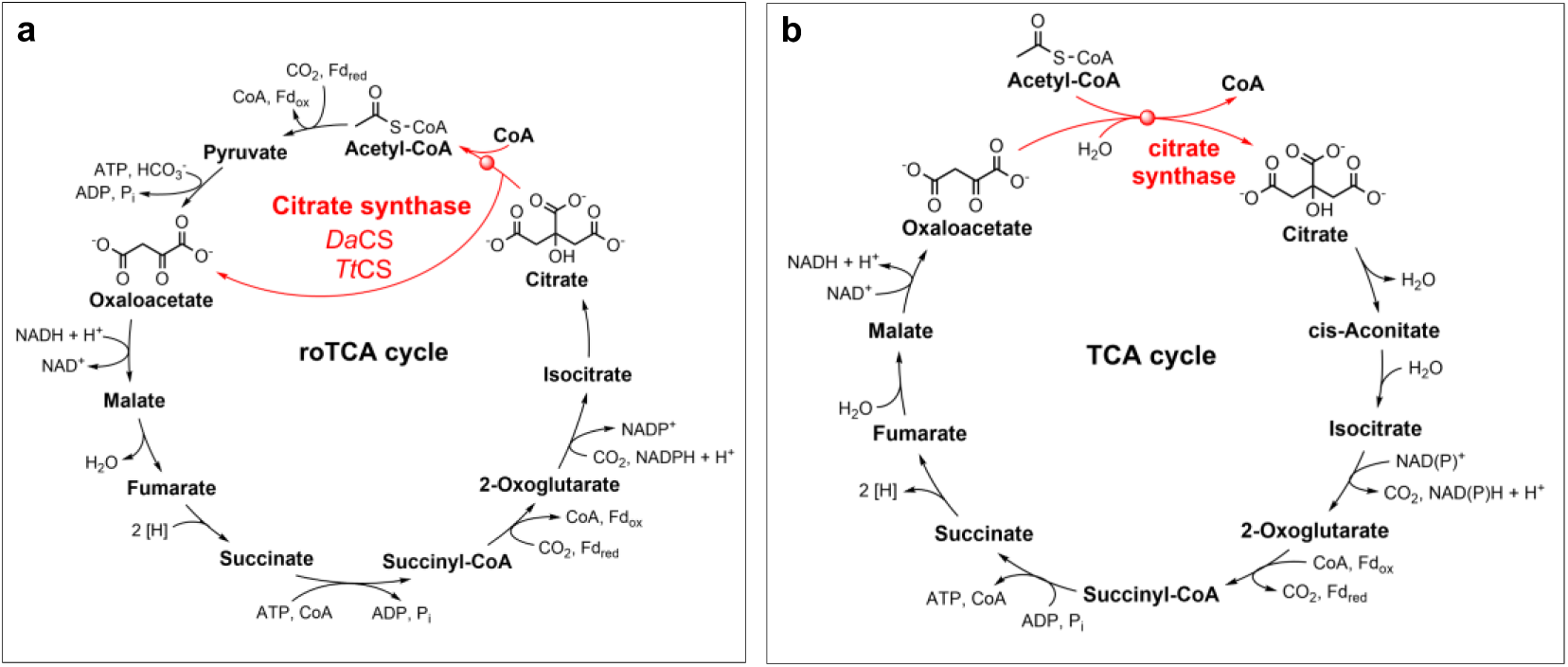
TCA cycle and reversed oxidative TCA cycle. **a,** Reversed oxidative tricarboxylic acid (roTCA) cycle, with citrate cleavage reaction shown in red. **b,** General oxidative tricarboxylic acid (TCA) cycle, with citrate synthesis reaction shown in red.

By reducing ATP consumption per round, roTCA cycle emerges as a more efficient autotrophic carbon fixation pathway^6^. Studies have shown that roTCA cycle is driven by high CO_2_ level^6^. High CO_2_ partial pressure promotes acetyl-coenzyme A carboxylation with CO_2_ to produce pyruvate, which is the next step of acetyl-CoA generation from citrate in roTCA cycle^6^. This leads to the equilibrium of reaction catalyzed by citrate synthase shift towards citrate cleavage^6^. As reported, citrate synthases of *D. acetivorans* (*Da*CS, Gene: Desace_08345)^6^ and *T. takaii* (*Tt*CS, Gene: TST_0783)^5^ catalyze citrate cleavage (Fig. 1) under thermodynamical challenge^5–6^ (free-energy change ΔG′ more than 35 kJ mol^−1^)^12^, but neither their catalytic mechanism nor directly regulatory factors have been identified.

As the forward reaction of citrate cleavage, a classical catalytic mechanism for citrate synthase had been proposed^9–10^, in which the first step of whole reaction is acetyl-group enolization of acetyl-CoA catalyzed by histidine and aspartate as general acid/base^13–14s^ (Supplementary Fig. 2b-c). The carbonyl carbon (C2) of oxaloacetate undergoes nucleophilic attack by the methylene of enolate^15^ and Claisen condensation happens between C2 and the enolated acetyl-group^14^, while (3*S*)-citryl-CoA is generated as intermediate (Supplementary Fig. 2d). The thioester bond of citryl-CoA hydrolyzes (Supplementary Fig. 2e) to produce citrate and coenzyme A (Supplementary Fig. 2f) which has no sufficient evidence and no plausible hypothesis^15–16^.

### Function of *Da*CS and *Tt*CS in normal air

High sequence identities (Supplementary Fig. 4, Supplementary Table 3-4) and similar kinetic properties of *Da*CS and *Tt*CS with citrate synthases from aerobic organisms^5–6^ suggest that the anaerobic environment needed by *D. acetivorans* and *T. takaii*^4–5^ may be not necessary for the function of their citrate synthases.

Citrate cleavage activities of *Da*CS and *Tt*CS were determined by ultra performance liquid chromatography (UPLC) ^4–6^ at different temperatures in normal air (Supplementary Fig. 5b). Citrate synthesis activities were also determined (Supplementary Fig. 5a) at same oxygen concentration. For *Da*CS, the amount of produced acetyl-CoA reached maximum equilibrium level at 70 ℃ along with increasing temperature (Supplementary Fig. 5b, Supplementary Fig. 5e), consistent with its increasing citrate synthesis activity (Supplementary Fig. 5a). As *Da*CS lost most of activity in forward reaction at 90 ℃, its acetyl-CoA production decreased sharply (Supplementary Fig. 5a-b). *Tt*CS demonstrated the highest production of acetyl-CoA and activity of citrate synthesis at 90 ℃ (Supplementary Fig. 5a-b, Supplementary Fig. 5f) which might be limited by detect method according to the trend in activity curve. Similar citrate cleavage activities were detected in four other citrate synthases from human (*h*CS), *Arabidopsis thaliana* (*At*CS), *Thermoplasma acidophilum* (*Tp*CS) and *Saccharolobus solfataricus* (*Sc*CS) which belong to eukaryotic mitochondria and thermophilic archaea respectively (Supplementary Fig. 3). The activity-temperature curves of these enzymes also show similarity with *Da*CS/*Tt*CS (Supplementary Fig. 5a-b).

The quantity of equilibrium reaction products generated by *Da*CS, *Tt*CS, or other four TCA citrate synthases progressively increase with rising temperature until protein denaturation occurs. This observation conforms to Le Chatelier’s principle that the extent of the thermodynamically unfavorable reverse reaction increases as temperature increasing, since it is an endothermic reaction (ΔH > 0) according to calculation. The total enthalpy change (Δ*H_m_*) of the reverse reaction catalyzed by citrate synthase was calculated by the following equation:

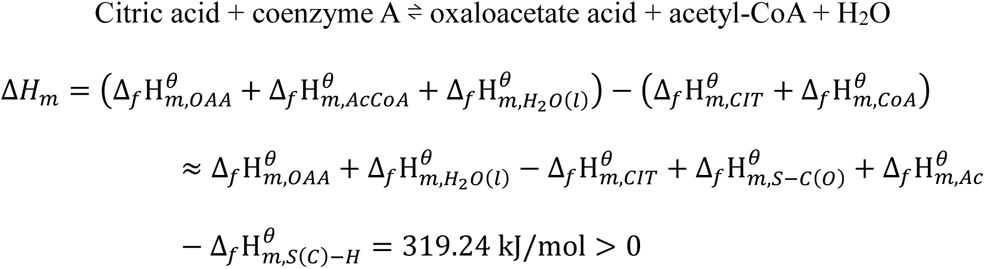

The standard molar enthalpy of formation for citric acid 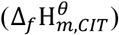 was −1543.8 kJ/mol^17^, for oxaloacetate acid 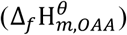 was −943.21 kJ/mol^18^, and for water 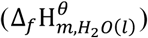 was −285.96 kJ/mol^19^. Because the standard molar enthalpies of formation for CoA compound were not available, we took Pauling’s assumption that the total bond enthalpy of an molecule is the sum of all its bonds^20^ to calculate the difference between the enthalpies of formation of CoA and acetyl-CoA using the standard molar enthalpies of formation for acetyl group 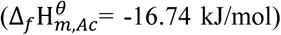, thiol bond 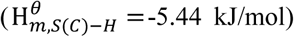^21–22^ and thioester bond 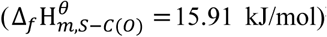^21–22^ with this equation: 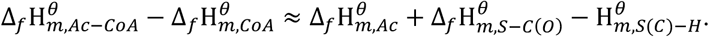

### Radical in reverse reaction

To confirm the chemical nature of the product, we provided [2, 4-^13^C] citrate and unlabeled coenzyme A (CoA) as substrates for *Da*CS to determine its citrate cleavage activity at 70 ℃ in air using nuclear magnetic resonance (NMR) spectroscopy. A clear peak at 30 ppm consistent with the chemical shift of [1-^13^C] acetyl-CoA (Fig. 2a, Supplementary Fig. 7a) shows its production, which corresponds to the UPLC results. To investigate the occurrence of reverse reaction under thermodynamically unfavorable conditions, electron paramagnetic resonance (EPR) was utilized to characterize substances involved in this process using 5,5-Dimethyl-1-pyrroline N-oxide (DMPO) as the spin trapping agent (Fig. 2, Supplementary Fig. 8). In the EPR spectrum of 70 ℃ heated *Da*CS reverse reaction mixture, a six-line signal corresponding to DMPO-alkyl radical adduct (A_N_ = 15.58 G, A ^β^ = 22.41 G, intensity ratio nearly equaling 1:1:1:1:1:1) and a four-line signal corresponding to DMPO-hydroxyl radical adduct (A_N_ =14.81G, A ^β^ = 14.81 G, intensity ratio ≈ 1:2:2:1) were identified (Fig. 2b, Supplementary Fig. 8e, Supplementary Table 1). No free radical was detected in unheated samples or sample containing substrates at much higher concentration (Fig. 2b, Supplementary Fig. 8f).

**Fig. 2:**
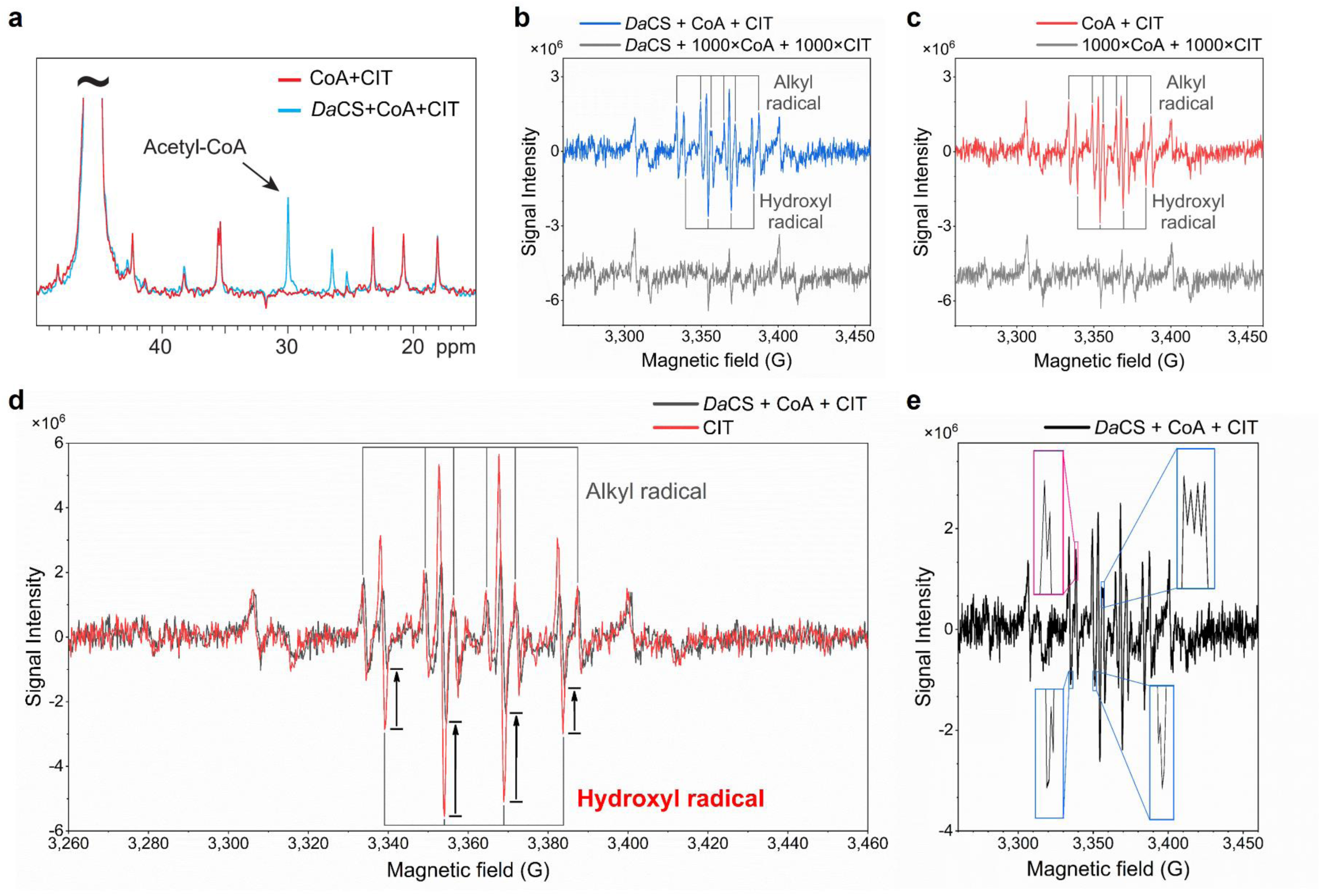
Acetyl-CoA and free radical production in *Da*CS catalyzed reverse reaction. **a**, ^13^C NMR analysis of [1-^13^C] acetyl-CoA production in the reverse reaction catalyzed by *Da*CS. **b**, EPR spectra of *Da*CS reverse reaction mixture containing 1 fold (10 μM) or 1,000 fold concentration (10 mM) CoA and citrate. In one fold group, six-line signal from DMPO-alkyl radical adduct (*A*_N_ = 15.58 G, *A*_H_^β^ = 22.41 G) and four-line signal from hydroxyl radical ones (*A*_N_ = 14.84 G, *A*_H_^β^ = 14.81 G) were identified. **c**, EPR spectra of 1 fold (10 μM) or 1,000 fold concentration (10 mM) CoA and citrate. In one fold group, signals from alkyl radical (*A*_N_ = 15.62 G, *A*_H_^β^ = 22.40 G) and hydroxyl radical (*A*_N_ = 14.86 G, *A*_H_^β^ = 14.85 G) were identified. **d**, Signal intensity of hydroxyl radical decreased in EPR spectrum of 10 μM citrate after adding *Da*CS and CoA. Signals from alkyl radical (*A*_N_ = 15.49 G, *A*_H_^β^ = 22.80 G) and hydroxyl radical (*A*_N_ = 14.84 G, *A*_H_^β^ = 14.81 G) were also detected in citrate. **e**, Enlarged split peak tips in EPR spectrum of *Da*CS-reaction sample. Peak tips from alkyl radical signal are shown in blue frame, while that from hydroxyl radical in pink.

In the EPR spectra of CoA contained sample, several abnormal tip splits were observed on peaks from both radical species (Fig. 2e, Supplementary Fig. 8g-i), and the intensity distributions of both peak groups deviated from corresponding theoretical values (Supplementary Table 1). Based on DMPO-thiyl radical adducts signals found in previous studies^23–24^ and the acetyl-CoA synthesis mechanism by pyruvate:ferredoxin oxidoreductase involving CoA-thyil radical^25^, we consider the tip-split peaks as the superposition of CoA-thyil radical and other radical species (Supplementary Table 1). In citrate spectrum, the intensity of hydroxyl radical signals surpassed that in sample containing CoA, while intensity of alkyl radical signals decreased slightly (Fig. 2d). These findings suggest that much more hydroxyl radical would be generated in heated citric acid solution compared with CoA, while a substantial portion of hydroxyl radical would be rapidly consumed by CoA or fixed by enzyme when mixed with them before trapped by DMPO.

To ascertain the function of radicals in reverse reaction, we analyzed *Da*CS reaction mixture with extra hydrogen peroxide (H_2_O_2_) by UPLC and observed that excess hydroxyl radicals had no impact on the reaction equilibrium (Supplementary Fig. 7d). Furthermore, the NMR spectra of *Da*CS reaction mixture with *L-*ascorbic acid or melatonin as free radical scavengers (Supplementary Fig. 7b-c) showed the unaltered equilibrium of reverse reaction. From the results, the enzyme prevent substance in solution from affecting radicals inside its catalytic pocket, including extra hydroxyl radical, free radical scavengers, and reductive citric acid.

The similarity between the spectra of mixed substrates with or without *Da*CS (Fig. 2b-c, Supplementary Fig. 8c, Supplementary Fig. 8e) indicate that all radicals detected are generated by heating citrate and CoA, but their lack of stability hindered interaction in the absence of enzymes. This was evidenced by NMR analysis, products were only observed in *Da*CS-containing substrates (Fig. 2a, Supplementary Fig. 8a).

According to these results, hydroxyl radicals act as catalysts in reverse reaction and alkyl radicals are more likely to exist in the enzyme’s catalytic pocket as transient intermediates. Enzyme catalytic pocket keeps a low concentration of substrates and protects the transient radicals from solvent environment.

### Structural basis of citrate cleavage from *Da*CS-intermediate complex

In *Da*CS-intermediate complex (PDB: 8XXC), the electron density in Chain E/F shows that *Da*CS is binding with an oxaloacetate and an acetate analogue (Fig. 3b-c, Supplementary Fig. 10c-f) cleaved from citrate, since neither oxaloacetate nor acetate analogues have been added in reservoir solution or crystal soaking solution. The ketone group of oxaloacetate forms hydrogen bonds with His^271^, Arg^328^, Arg^400^ and its carboxyl group form hydrogen bonds with His^235^, His^319^ and Arg^420*^ as in *Da*CS-citrate complex (Fig. 3a-c). The acetate analogue in catalysis pocket is stabilized by a hydrogen bond network formed by active-site residues and nearby water molecules, with different conformation in Chain E and Chain F (Fig. 3b-c).

**Fig. 3:**
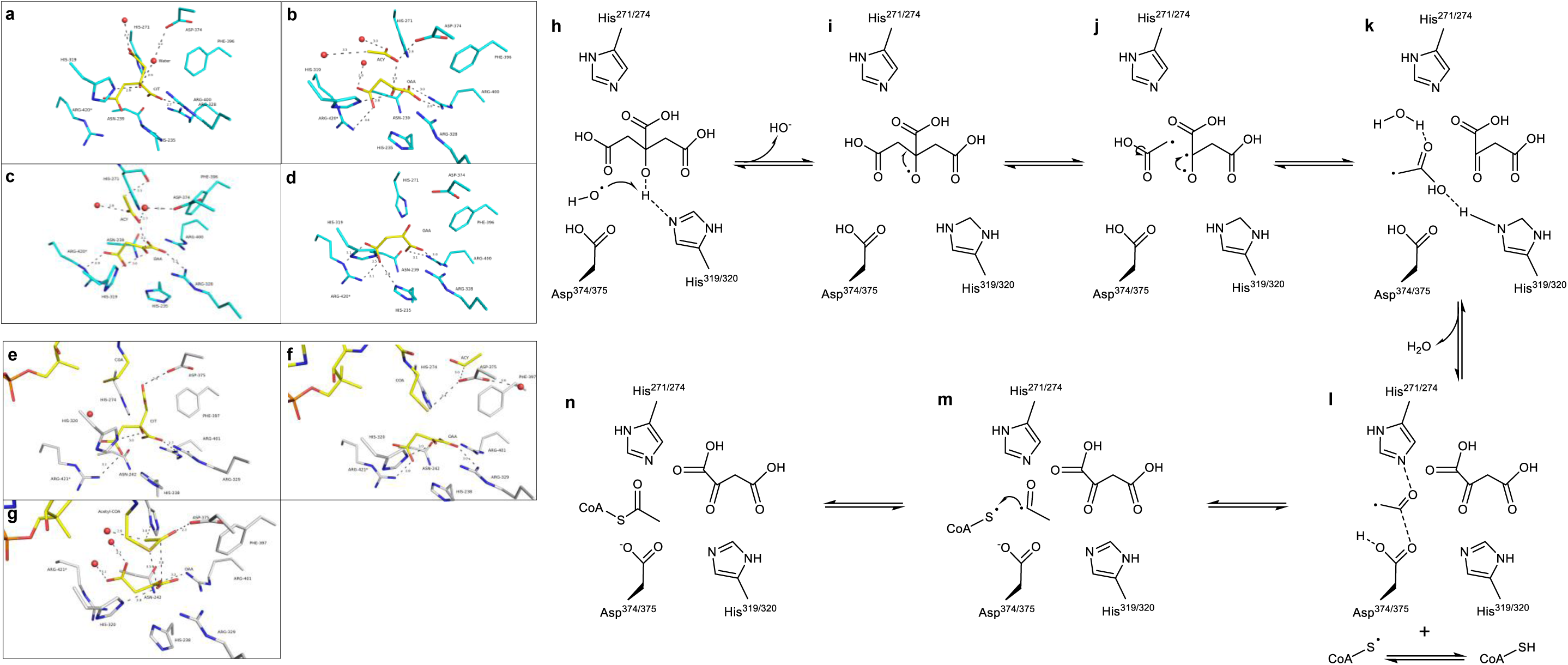
Structures of intermediate in active site of *Da*CS and *Tt*CS, and proposed mechanism of citrate synthase reverse reaction. **a**, Interactions (dashes) of citrate (yellow) to *Da*CS active site residues (cyan) and water (red), from *Da*CS-citrate complex (PDB: 8XU2). **b**, Interactions of oxaloacetate and acetate analogue as intermediate in *Da*CS active site, from Chain E in *Da*CS-intermediate complex (PDB: 8XXC). **c**, Interactions of oxaloacetate and acetate analogue of another conformation, from Chain F in *Da*CS-intermediate complex (PDB: 8XXC). **d**, Interactions of oxaloacetate, from *Da*CS-oxaloacetate complex (PDB: 8XU1). **e**, Interactions (dashes) of citrate and coenzyme A (yellow) to *Tt*CS active site residues (gray) and water (red), from *Tt*CS-citrate/CoA complex (PDB: 8XXD). **f**, Interactions of oxaloacetate, intermediate acetyl group, and CoA, from Chain C in *Tt*CS-intermediate complex (PDB: 8XXE). **g**, Interactions of oxaloacetate and acetyl-CoA, from Chain F in *TtCS*-oxaloacetate/acetyl-CoA complex (PDB: 8XXF). **h-n**, Proposed mechanism of citrate synthase reverse reaction. Unpaired electron represents free radical. The residue numbers come from *Da*CS/*Tt*CS.

In Chain E, methyl of acetate analogue towards to oxaloacetate and is stabilized by hydrogen bond network formed by Leu^270^, His^271^, His^319^, Asn^372^, Asp^374^ and several water molecules (Fig. 3b). The electron density suggests that the binding of acetate analogue in the catalytic pocket is transient and its methyl rotates 90 degrees away from oxaloacetate (Fig.3b, Supplementary Fig. 10c-d). In Chain F, the acetate analogue is stabilized by another hydrogen bond network consisting of Leu^270^, His^271^, Asn^372^, Asp^374^, oxaloacetate and water molecules (Fig. 3c). The methyl of acetate analogue rotates 180 degrees away from oxaloacetate and forms hydrogen bond with His^271^, while two oxygen atoms of acetate analogue form hydrogen bonds between two water molecules nearby (Fig. 3c).

### Structural basis of CoA acetylization from *Tt*CS-intermediate complex

In Chain C of *Tt*CS-intermediate complex (PDB: 8XXE), the density, observed near the catalytic residue Asp^375^ (Asp^374^ in *Da*CS) and substrate CoA, is suspected as an acetyl group (Fig. 3e, Supplementary Fig. 11c-d). The suspected acetyl group is stabilized in catalytic pocket by a hydrogen bond network consisting of His^274^, Ala^277^, Asp^375^ (mainchain O of His^274^, mainchain N of Ala^277^ and sidechain of Asp^375^) and water molecules (Fig. 3f). Ketone group of oxaloacetate forms hydrogen bonds to His^272^, Arg^329^, Arg^401^ and its carboxyl groups form hydrogen bonds with His^236^, His^320^ and Arg^421*^ (Fig. 3f). The binding of CoA resembles that in *Tt*CS-citrate/CoA complex, binding with loop 314-321 and loop 365-369 (Supplementary Fig. 12a-d). In Chain B, the ketone group of oxaloacetate flips 180 degrees away from Arg^421^, and the binding position of CoA changes to loop 39-44 and loop 314-321 closer to cleft entrance (Supplementary Fig. 13a). This conformational change is accompanied by the movement of original CoA binding loop, loop 314-321 and loop 365-369 (4.79 Å and 3.93 Å), which widens the cleft for CoA entry.

In Chain F of *Tt*CS-oxaloacetate/acetyl-CoA complex (PDB: 8XXF), the produced acetyl-CoA adopts the binding form almost identical to CoA in Chain D of *Tt*CS-citrate/CoA complex while its acetyl group forms hydrogen bonds with His^274^ and Asp^375^ (Fig. 3g). Oxaloacetate in *Tt*CS-oxaloacetate/acetyl-CoA complex rotates 180 degrees from its position in *Tt*CS-intermediate, and the two carboxyl oxygens of ketone group form hydrogen bonds with sidechain of Arg^401^ (Fig. 3g). Because of this rotation, the distance between oxaloacetate and acetyl group of acetyl-CoA comes to 3.8 Å which may prevent reverse citrate synthesis (Fig. 3g). Acetyl-CoA shows another binding form in Chain A of *Tt*CS-oxaloacetate/acetyl-CoA complex, loop 39-44, loop 314-321 and Met^416*^-Ile^419*^ form the new acetyl-CoA binding site when loop 365-369 is disordered, which indicates the release process of acetyl-CoA (Supplementary Fig. 13b).

### Mechanism of reverse reaction

Based on abundant hydroxyl radical detected by EPR in heated citric acid solution and several water molecules or hydroxyl molecules near citrate in *Da*CS-citrate complex (PDB: 8XU2) (Fig. 3a), we consider that hydroxyl radicals participate in citrate cleavage and play the role of energy donors. Consequently, we present a mechanistic model for the reverse reaction.

From the structure of *Da*CS-intermediate, there is a water molecule considered as hydroxyl radical from EPR result (Fig. 2b) located near the hydroxyl oxygen of citric acid (2.76 Å) (Fig. 3a, Supplementary Fig. 10a-b). After citric acid binding, the hydroxyl radical starts the reaction by binding nearby the hydroxyl of citric acid and transfer the radical electron to its hydroxyl oxygen, while the imidazole nitrogen of His^319^ receives and transfers the hydroxyl hydrogen atom (Fig. 3h). The activated oxygen atom with an unpaired electron attacks C3 atom of citric acid, leading to the cleavage of the C2-C3 bond and generating an acetic acid analogue having an unpaired electron (Fig. 3i-j) which corresponds to the alkyl radical identified in the EPR analysis (Fig. 2b). At this stage, activated oxygen atom forms carbon-oxygen double bond with the C3 atom and oxaloacetic acid is produced as the first product (Fig. 3j-k). After cleavage, methyl of produced acetic acid analogue rotates 180 degrees in the catalytic pocket (Fig. 3c, Fig. 3j-k), preparing for next state. Throughout this process, the acetic acid analogue exists as monohydrate for stability and translation (Fig.3b-c, Fig. 3k). From *Tt*CS-intermediate structure, the acetic acid radical dehydrates to form acetyl radical, leaving an acetyl radical stabled by His^274^ and Asp^375^ (Fig. 3f, Fig. 3k-l). After the water molecule is released, the acetyl radical approaches the thiol group of CoA assisted by His^274^ and Asp^375^ (Fig. 3m). From the acetyl-CoA synthesis mechanism of pyruvate:ferredoxin oxidoreductase^25^ and the split peak tips in EPR results of samples containing CoA (Fig. 2e, Supplementary Fig. 8g-i), it suggests that CoA generated thyil radical at high temperature (Fig. 3l). In the final process of the reverse reaction, the CoA-thyil radical forms a covalent bond with the acetyl radical in the catalytic pocket and produces acetyl-CoA (Fig. 3m-n).

From EPR results, the hydroxyl radical generated by substrates may be far excessive for the reaction (Fig. 2d). And the probable radical rich environment formed in the enzyme catalytic pocket is also considered as key factor for the reaction to overcome the energy barrier, although there is only one hydroxyl radical involved in the reaction. We hypothesize that the thermodynamically unfavorable reverse reaction obtains a portion of its required energy from high temperature environment. High temperature not only provides energy and increases free radical production in solution, but also promotes molecular motion and flexibility of enzymes, as evidenced by the increasing extent of reverse reaction observed across all citrate synthases with rising temperature (Supplementary Fig. 5b).

The involvement of aerobic environment in the catalytic mechanism mentioned above is negligible, as indicated by the lack of contribution to activity of *Da*CS and *Tt*CS, two citrate synthases involved in the roTCA cycle, which is supported by experiments conducted under normal air conditions (Supplementary Table 5). The results also support with the opinion of Steffens, L. et al. that high CO_2_ partial pressures accelerate the removal of produced acetyl-CoA by pyruvate synthase^6^, rather than directly influencing the catalysis of citrate synthase.

### Peroxide bond in human citrate synthesis

Since the unusual reaction mechanism of citrate cleavage in roTCA cycle and unclarified details remained in citrate synthesis mechanism^15–16^, we use Raman spectroscopy and X-ray diffraction with time-resolved character to capture the intermediate states in human citrate synthase (*h*CS). Three characteristic bands falling within the expected range for ν(O-O) frequencies^26–29^ were detected in Raman spectra from different *h*CS samples.

In spectra A, the band at 745 cm^-1^ appears after 300 sec reaction which is unclear in spectra of that reacting 120s or 440s (Fig. 4a). The same reaction time is used in *h*CS-intermediate 2 (PDB: 8ZVR) collection. Electron density observed near CoA in the structure (Fig. 4d) can be matched as acetyl group and oxaloacetate rather than citrate (Supplementary Fig. 16d-f, Supplementary Fig. 18). The distance between acetyl oxygen (O1^(ACE)^, Fig. 4d) and ketone oxygen of oxaloacetate (O3^(OAA)^, Fig. 4d) is 1.4 Å, while the distance between -CH_3_ of acetyl group (C2^(ACE)^, Fig. 4d) and ketone carbon of oxaloacetate (C3^(OAA)^, Figure 1D) is 2.5 Å (Fig. 4d).

**Fig. 4:**
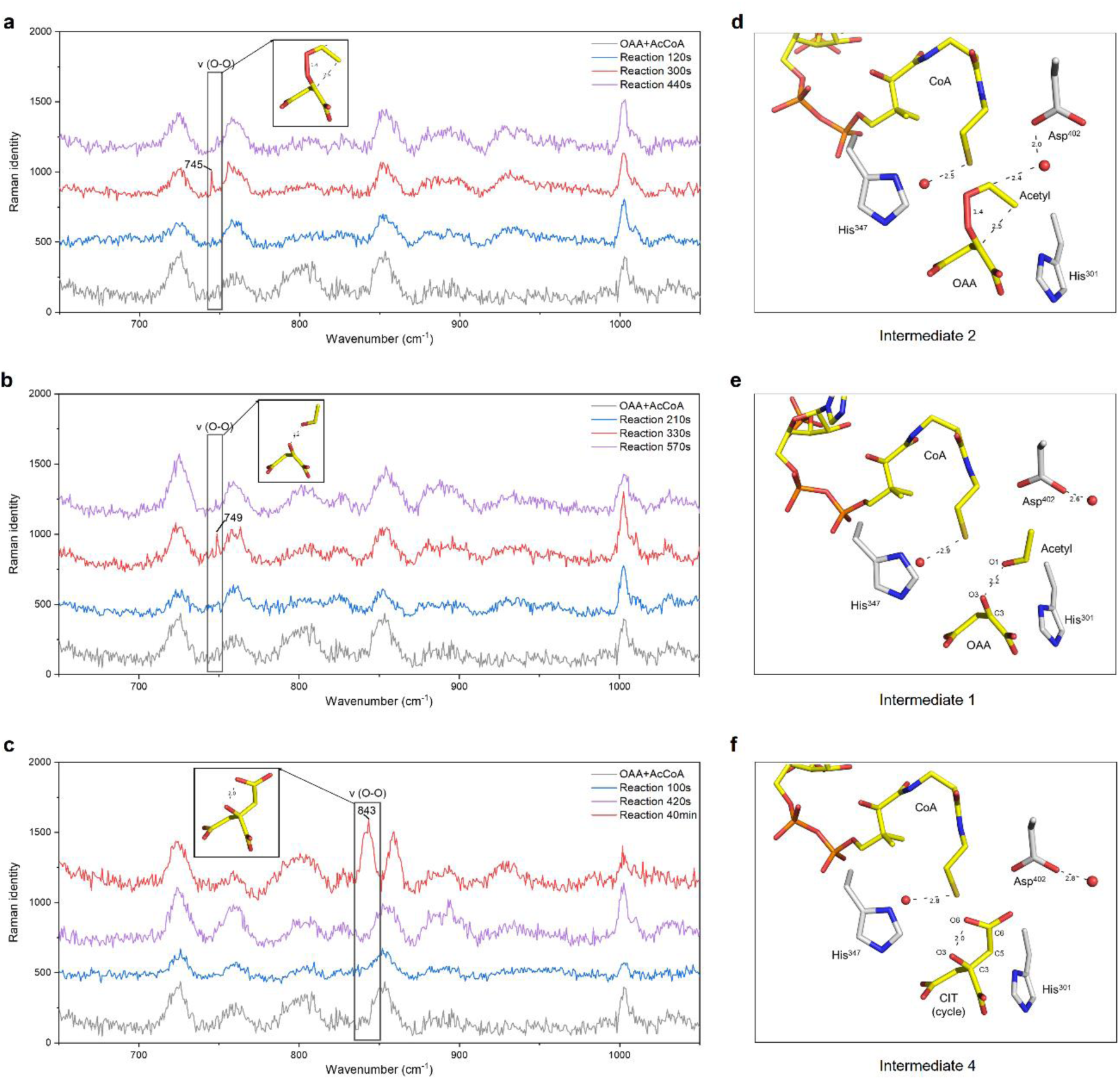
Raman spectra of *h*CS crystal range of 650-1050 cm^-1^ for different reaction times. **a,** Raman spectra of *h*CS crystal after mixed with substrates for 120s (blue), 300s (red) and 440 (purple). The concerned band appears at 745 cm^-1^ in spectrum of 300s and labeled. The corresponding intermediate structure are shown in yellow sticks. Raman spectra of oxaloacetate and acetyl-CoA is showing in gray. **b,** Raman spectra of another *h*CS crystal at reaction time = 210s (blue), 330s (red) and 570s (purple). The concerned band appears at 749 cm^-1^ in spectrum of 330s. The corresponding intermediate structure are shown in yellow sticks. Raman spectra of oxaloacetate and acetyl-CoA is showing in gray. **c,** Raman spectra of another *h*CS crystal at reaction time = 100s (blue), 420s (purple) and 40 min (red). The concerned band appears at 843 cm^-1^ in spectrum of 40 min. The corresponding intermediate structure are shown in yellow sticks. Raman spectra of oxaloacetate and acetyl-CoA is showing in gray. **d,** *h*CS-intermediate 2 structure (PDB: 8ZVR). Ligand is colored in yellow with atoms labeled and distances annotated in dash. Proteins colored in gray. **e,** *h*CS-intermediate 1 structure (PDB: 8ZVM). **f,** *h*CS-intermediate 4 structure (PDB: 8ZVU).

The interaction between the two oxygen atoms is suspected as peroxide bond according to spectra^27–28^ and structure, which also indicates no C-C bond between C2^(ACE)^ and C3^(OAA)^ (Supplementary Fig. 16d-f).

Another band at 749 cm^-1^ appears in spectra B with 330 sec reaction time which is unclear in that reacting 210s or 570s (Fig. 4d). Similar shifts of bands in spectra A and B indicate similar O-O interactions. The corresponding intermediate of 749 cm^-1^ band in spectra B is observed in *h*CS-intermediate 1 structure (PDB: 8ZVM), in which O1^(ACE)^ (Fig. 4e) has 2.2 Å interaction with O3^(OAA)^ (Fig. 4e, Supplementary Fig. 16a-c). The water molecular nearby Asp^402^ contributes to catalyzing the hydrolysis of thioester bond in acetyl-CoA (Fig. 4e, Supplementary Fig. 14a-c).

The band at 843 cm^-1^ appears in spectra C after 40 min reaction and disappears in these reacting shorter times (Fig. 4c). The strong band represents stable O-O bond^28–31^ which indicates reaction equilibrium, and the corresponding structure is matched in *h*CS-isntermediate 4 (PDB: 8ZVU). The O3, C3, C5, C6 and O6 of citrate analogue (Fig. 4f) exhibit coplanar and form five-membered ring according to the electron density (Supplementary Fig. 16j-l), indicating a peroxide bond in equilibrium.

Another intermediate state obtained (*h*CS-intermediate 3, PDB: 8ZVT) shows acetate analogue and oxaloacetate binding in active site (Supplementary Fig. 16g-i, Supplementary Fig. 19) with 1.9 Å distance between O1^(ACY)^/O3^(OAA)^ and 2.1 Å distance between C2^(ACY)^/C3^(OAA)^ (Supplementary Fig. 14e), which indicate the formation of O-O bond and the generation of C-C bond. Four additional structures obtained represent initial states and final sates (PDB: 8ZVL, 8ZW1, 8ZVV, 8ZVW). In *h*CS citrate complex (PDB: 8ZVV), the C6-carboxyl of citrate are not forming plane with O3, C3, C5 like *h*CS-intermediate 4 and all the atoms of citrate obey stereochemical constrains without strong interaction between O3 and O6 (Supplementary Fig. 17a-c) according to the 2.9 Å distance (Supplementary Fig. 14g).

### Concurrent multi-mechanisms in citrate synthesis process

Intermediates indicate multiple concurrent citrate synthesis mechanisms in citrate synthesis (Fig. 5). In mechanism I and II, the ketone oxygen of acetyl group from acetyl-CoA have interaction with the ketone oxygen of oxaloacetate (O3, Supplementary Fig. 14b) to form peroxide bond. The formation of peroxide bond leads to the production of two unpaired electrons between the bonds of CH_3_-C-O (O1, C1, C2, Supplementary Fig. 14d) and C=O (C3, O3, Supplementary Fig. 14d). The unpaired electron on acetyl group is delocalized in O1-C1-C2 bond, promoting one methyl proton to transfer to His^347^ (Fig. 5). Mechanism I and II have differences in whether C-C bond generation or thioester bond in acetyl-CoA hydrolysis occurs first.

**Fig. 5:**
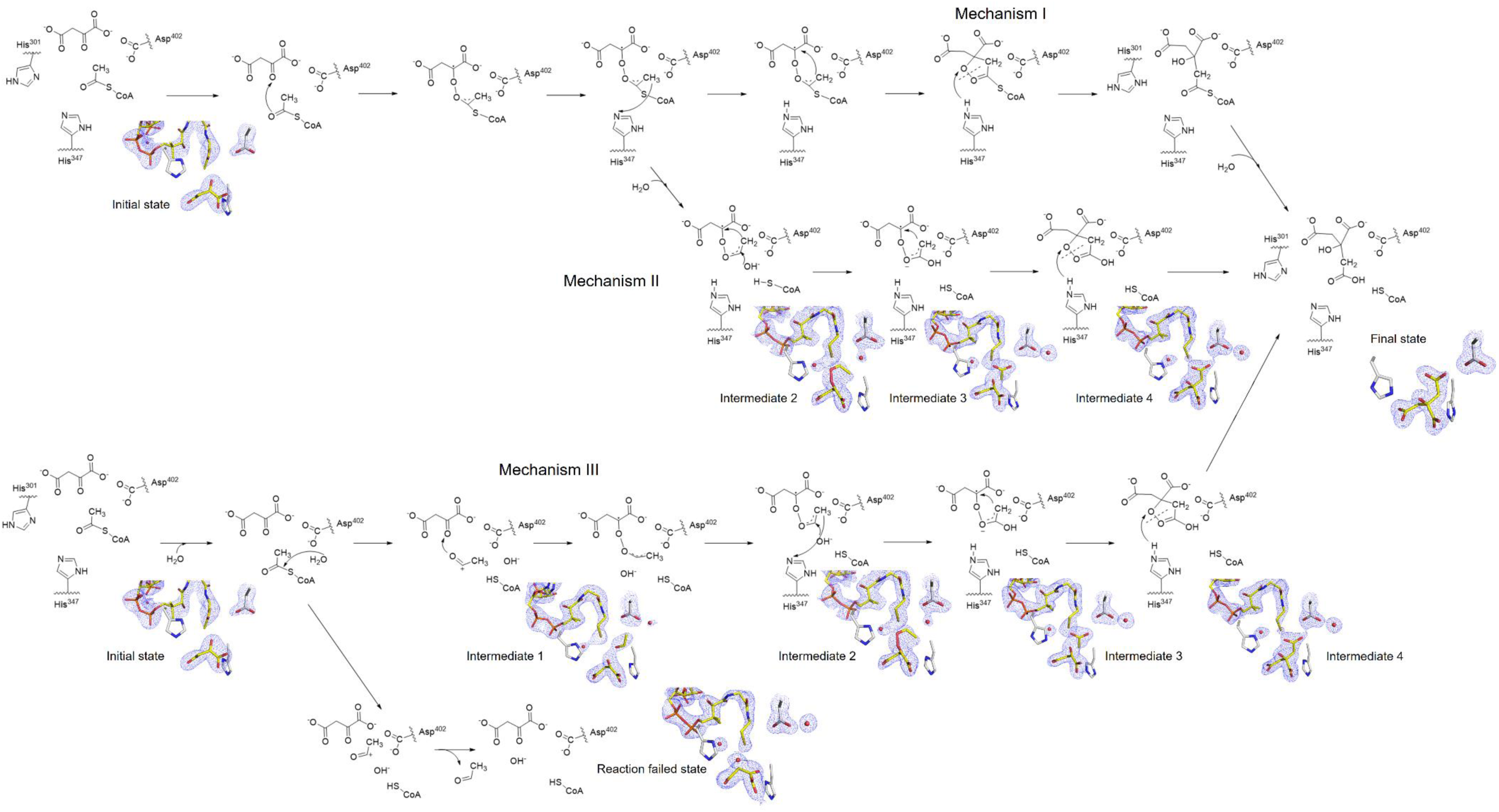
Multi catalytic process of *h*CS. Mechanism I and II shows the catalytic mechanism centered on citryl-CoA, and they share the first four steps. Mechanism III describe the catalytic process of acetyl-CoA direct hydrolysis without forming citryl-CoA, and it shares the first two steps with failed process. The corresponding intermediate structures are shown nearby.

In Mechanism I, the two unpaired electrons on C3^(OAA)^ and C2^(ACE)^ promote the five-membered ring intermediate generation to form C-C bond. The peroxide bond is then broken and citryl-CoA is generated as intermediate with hydroxyl re-taking the proton on His^347^ ^15^. The citryl-CoA is hydrolyzed according to classic mechanism^15^ (Supplementary Fig. S2e), producing citrate and CoA (Fig. 5). In Mechanism II, the thioester bond of acetyl-CoA is hydrolyzed, while the generated acetyl group is linked to oxaloacetate by peroxo-bridge. The acetyl group undergoes nucleophilic attack by the hydroxyl group nearby and transforms into acetate analogue. C2^(ACE)^ and C3^(OAA)^ (Supplementary Fig. 14d) form C-C bond by the unpaired electrons and generate citrate analogue intermediate containing five-membered ring with peroxide bond between O6^(CIT)^ and O3^(CIT)^ (Supplementary Fig. 14f). This intermediate turns into citrate with free conformation in the active site after the peroxide bond broken (Fig. 5, Supplementary Fig. 14g).

In mechanism III, the thioester bond of acetyl-CoA is firstly hydrolyzed by water molecular nearby CoA, generating free acetyl group with positively-charged ketone carbon. The hydroxyl (hydroxide) from water molecular remains in the active site stabled by Asp^402^ and CoA. The O1^(ACE)^ forms peroxide bond with O3^(OAA)^ (Supplementary Fig. 14d). The acetyl group is hydroxylated and forms an acetate group, while one proton on its methyl is transferred to His^347^. C2^(ACY)^ and C3^(OAA)^ (Supplementary Fig. 14e) form C-C bond and generate five-membered ring citrate intermediate. The peroxide bridge bond is then broken, and the hydroxyl group gets the proton from His^347^, transforming the intermediate into citrate (Fig. 5).

In the whole catalytic process, it is possible that catalysis is failed according to the structural results, and it shares the first two steps with mechanism III while the acetyl group produced by acetyl-CoA hydrolysis fails to form peroxide bond with oxaloacetate, and the hydroxyl group produced from hydrolysis is attracted by oxaloacetate (Fig. 5, Supplementary Fig. 14h).

### Peroxo bridge like interaction promotes C-C bond generation by cyclization

Orbital hybridization theory provides an explanation for how the peroxide bond and five-membered ring promotes the generation of C-C bond which act as peroxo bridge. Both the ketone carbon of acetyl group (C1) and the ketone carbon of oxaloacetate (C3) is sp^2^ hybrid (Fig. 6a). Peroxide bond formation breaks both C=O double bonds while the carbon atoms (C1, C3) are still in sp^2^ hybrid with half-empty p-orbitals between C-O (Fig. 6b). The hybridization and p-orbitals in C3-O3-O1-C1 form conjugated connection similar to conjugated double bonds, and promote the coplanarity of five atoms while the three atoms of acetyl/acetate analogue (O1, C1, C2) are coplanar (Fig. 6b). The 120 °angle of O1-C1-C2 in acetyl/acetate analogue, the 110 °angel of O3-O1-C1 (Fig. 4d) and the 139 °angel of C3-O3-O1 (Fig. 4d) promote C2^(ACY)^ and C3^(OAA)^ in suitable conformation for cyclization. Driven by the two conjugated unpaired electrons, C2-C3 bond is generated and the cyclization reaction of C3-O3-O1-C1-C2 is done (Fig. 6c). Meanwhile, C-C bond generation causes the hybrid orbitals C3^(OAA)^ change from sp^2^ to sp^3^ (Fig. 6c) which breaks the conjugated connection and changes the bond angle, while the C2^(ACY)^ is sp^3^ hybrid originally (Fig. 6c). The change of conjugated orbital and bond angle breaks the five-atoms-coplane, weakening the peroxide bond, which would be broken naturally after leaving the catalytic environment (Fig. 6d). The peroxide bond generation is catalyzed by the residues in active pocket, and the formation of coplanar ring reduces the energy barrier of C-C bond generation.

**Fig. 6:**
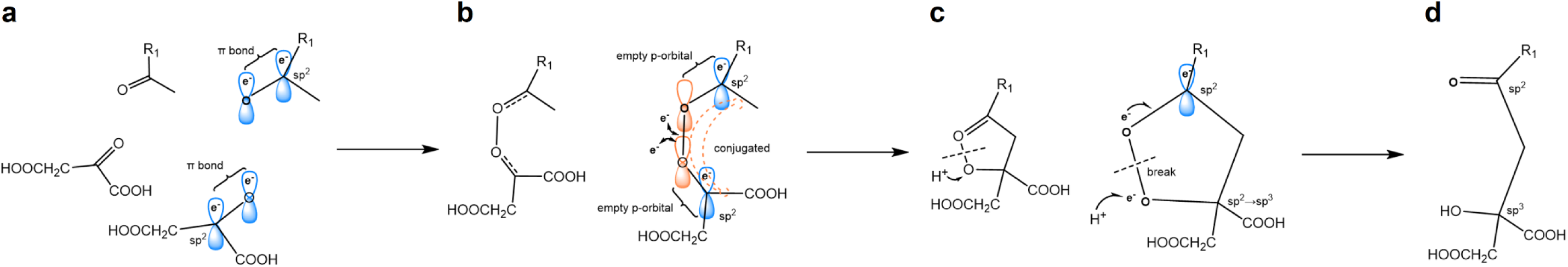
Orbital hybridization of peroxide bond, cyclization and C-C bond generation. **a,** Hybrid orbitals of reaction start state. **b,** Hybrid orbitals of peroxide bond formation and conjugated double bonds-like connection formation. **c,** Hybrid orbitals of five-membered ring cyclization formation and break. **d,** Final state of reaction. e^-^ represents electron. Hybrid orbitals colored in blue are with electron in it, colored in orange are empty. R_1_=-H, -OH, or -S-CoA.

### Peroxo bridge like interaction in C-C bond generation and cleavage

The synthesis and cleavage of citrate catalyzed by citrate synthase rely on the formation and break of the interaction between oxygen atoms. In citrate synthesis, a peroxo bridge like interaction is formed between carboxyl oxygen atom and ketone oxygen atom. The electrostatic field and intermolecular interaction network consist of amino acid residues and CoA in enzyme catalytic pocket contribute to the peroxo bridge like O-O bond formation. The O-O interaction, while the electron of the two oxygen atoms is paired to form O-O bond, breaks the two carbon-oxygen double bonds, and the unpaired-electron of carbon atoms retained in the p orbitals of broken C=O bonds which presented as half-empty orbitals. The empty p orbitals retained make the four connected atoms and the methyl carbon atoms of the acetyl group co-plane and form a connection similar to the conjugated double bond, which allow the unpaired electrons of carbon atoms in a delocalized conjugated state. The conjugation of electrons makes the methyl carbon atoms of the acetyl group or acetate group active and lose hydrogen, which make C-O-O-C-C cyclization to form carbon-carbon bond (C3-C2) with the influence of the co-plane and bond angle.

The empty p orbitals provide additional orbitals for the bonding electrons of O-O, which are in superposition state of multiple states. The stereochemical constraints of the cyclized atoms make the interaction between oxygen and oxygen extremely unstable, and the distance between the atoms varies from 1.3 Å for covalent bonding to 2.1 Å for hydrogen bonding. According to the observations, the oxygen-oxygen interaction may collapse into two types of peroxide bond and oxygen-oxygen hydrogen bond, depending on whether the electrons are in the bonding orbital or the empty p orbital.

The sp^3^ orbital of carbon atoms, forming along with C-C bond formation, and the paired electron in sp^3^ orbital break the conjugated double bond like state of the five-membered ring, which weaken the O-O interaction by stereochemical constraint. The bonded electron in O-O bond tend to the empty p orbitals of oxygen atom from the superposition state, and form a carboxyl group and a negatively charged oxygen atom. Due to the joint constraints of the catalytic pocket and coenzyme A, the distance between the two oxygen atoms is still less than 2.3 Å, and the returning electrons of the two oxygen atoms are still in a state of bonding and are still in a multiple superposition state. The structure of the enzyme changes after CoA release weakens the external stereochemical constraint, which break the unstable O-O interaction. The paired electron in O-O bond completely collapse into the oxygen atom p orbital state, and hydrogen bind negatively charged oxygen atom, citrate is formed.

In the reaction of citrate cleavage to form acetyl-CoA, the C-C bond between the methylene carbon of citrate and the keto carbon itself rotates with the increase of ambient temperature. The molecular thermal motion is intensified and the rotation of carboxyl group to the direction of keto hydroxyl group increases along with ambient temperature. Meanwhile, the environmental energy enhances the activity of electrons on oxygen atom and the possibility of hydrogen atom dissociation on keto hydroxyl group, which promotes the formation of peroxide bond like oxygen-oxygen interaction. After the oxygen-oxygen interaction is formed, the carboxyl oxygen atom remains an empty p orbital and the bonding electrons have the possibility of multiple superposition states and conjugation in the five-membered ring. The stereochemical constraint changes of the five atoms and the destruction of superposition electrons on the oxygen atom weaken the covalent bond between the five atoms. According to the different stereochemical constraints of each atom and the position of superposition electrons, the C-C bond between two carbon atoms with sp^3^ hybridization in the five-membered ring and the adjacent C-O bond are considered to be easy to break. The breaking of the C-C bond and C-O bond destroy the weak co-plane state of the five-membered ring, which also leads to the breakdown of the oxygen-oxygen interaction. The destroyed C-C bond forms alkyl radical and the charged oxygen atom come from oxygen-oxygen interaction forms hydroxyl radical with free electron, which correspond to the free radical experiment in the reverse reaction results.

When citrate enters the enzyme catalytic pocket, CoA and the catalytic environment have more strictly stereochemical constraint to fix the conformation of carboxyl group of citric acid toward the ketone hydroxyl group, making the above C-C/C-O bond break with greater probability of occurrence. The nearby histidine in catalytic pocket provides hydrogen, which reduces the possibility of carbon-oxygen bond breaking and increases the proportion of carbon-carbon bond breaking. With the help of enzymes, the reaction process develops to greater extent to produce acetic acid free radicals, and further reacts with CoA to produce acetyl-CoA.

### Conserved reaction in the citrate synthases family

Based on our verification of the reverse activity of other citrate synthases in the TCA cycle (Supplementary Fig. 5-6) and the highly conserved core structures among citrate synthases from various organisms (Supplementary Table 3-4), we propose that the mechanism of citrate cleavage and generation are widely present in the citrate synthase family across archaea to mammals, both anaerobic and aerobic. However, the distribution of citrate cleavage mechanism is restricted due to its requirement for high temperature environments and the negative impact on cells from free radicals involved. While citrate synthases from thermophilic archaea adopted thermostable structural features for adapting extreme environment^32–33^, the evolutionary-distant eukaryotic citrate synthases chose flexibility for high efficiency at lower temperature (Supplementary Fig. 5a). Both types of citrate synthases retain their ability to catalyze the reverse reaction despite lacking physiological significance. Although *Da*CS and *Tt*CS do catalyze citrate cleavage in roTCA cycle - one of the oldest metabolic pathways - they exhibit structures more similar to eukaryotic citrate synthases, suggesting a potential common origin with those found in eukaryote mitochondria while incorporating a few thermostable features.

These findings also provide circumstantial evidence supporting the endosymbiont hypothesis of mitochondrion. In the mitochondrion of eukaryote, its radical-rich environment allows the maintenance of multifunctional citrate synthases. Considering that reversible citrate synthesis occurs within its radical-rich matrix during TCA cycle operation, the reversible citrate synthases in mitochondrion may imply the potential possibility of roTCA cycle there.

In addition, the multiple concurrent mechanism in citrate synthesis process rather than the process with citryl-CoA intermediate we proposed explain how peroxo bridge like interaction promotes the C-C bond generation and cleavage by cyclization. Our findings also optimize the classical catalytic mechanism for citrate synthase in TCA/roTCA cycle with concise and adaptable time-resolved characteristic experimental approaches.

## Data availability

All data are available in main text or supplementary materials and from the Protein Data Bank (www.rcsb.org/).

## Acknowledgements

This project is supported by National Key Research and Development Program of China (Grant Nos. 2019YFA0904100) and National Natural Science Foundation of China (Grant Nos. T2221005). We sincerely thank the staff of Shanghai Synchrotron Radiation Facility, instrument BL18U1 and BL10U2 for the support in diffraction data collection. We appreciate L. Yu for EPR experiments.

## Author contributions

Y-J.F. prepared the protein samples and conducted the enzyme activity analysis. L.Y. and Y-J. F. conducted the X-ray experiments and analyzed the experimental data. Y-J. F. conducted the Raman spectrum experiments and analyzed the data. L.Y. and Y-J.F. conducted the NMR and EPR experiments and analyzed the experimental data. L.Y., Y-J.F. and W-M. G. wrote the manuscript. All authors read and approved the paper.

## Additional information

**Supplementary information** Supplementary Information is available for this paper.

**Correspondence and requests for materials** should be addressed to Weimin Gong.

## Peer review information

**Reprints and permissions information** is available at http://www.nature.com/reprints.

