## Supplemental figure 1-25, Supplemental table 1-6 for "C-C bond cleavage and generation in TCA cycle driven by radicals and peroxo-bridge like interaction"

### Materials and methods

#### Expression and purification of citrate synthases

The amino acid sequence of Desace_08345 from *Desulfurella acetivorans* (*Da*CS) was cited from the result of mass spectrometry described by Steffen, L. et al.[1], while the sequences of other five citrate synthases from human (*h*CS), *Arabidopsis thaliana* (*At*CS), *Saccharolobus solfataricus* (*Sc*CS), *Thermoplasma acidophilumcitrate* (*Tp*CS) and *Thermosulfidibacter takaii* (*Tt*CS) were from **Uniprot** (www.uniprot.org).

All sequences were codon-optimized and subcloned into the pET22b vector without signal peptides (for *h*CS and *At*CS), then were expressed in *E. coli* BL21 (DE3). The cells were grown at 37 °C in Luria-Bertani medium containing 50 mg/L ampicillin till the culture's $OD_{600\ nm}$ reached 0.6 to 0.8. Then expression was induced by 0.5 mM isopropyl-β-D-thiogalactopyranoside (IPTG) at 16 °C for 16~18 hours.

Cells containing recombinant *Da*CS were collected, resuspended in 20 mM Tris-HCl (pH 8.0), 100 mM NaCl and 20 mM imidazole and lysed by high pressure (80 MPa). Cell extracts were heated to 65 °C for 10 min[2]. After centrifugation, the supernatants were loaded on Ni-NTA column (GE healthcare) equilibrated with 20 mM Tris-HCl (pH 8.0), 100 mM NaCl containing 20 mM imidazole. The column was washed with the same buffer containing 20 mM and 50 mM imidazole and then protein was eluted with buffer containing 500 mM imidazole[1]. *Da*CS was concentrated after imidazole was removed using Amicon Ultra-15 mL 30 kDa centrifugal filters (Milipole). For *Da*CS used in NMR or EPR measurements, TRIS was removed as well as imidazole with 50 mM $Na_2HPO_4$-$KH_2PO_4$ (pH 7.0) and 100 mM NaCl.

Recombinant *Sc*CS and *Tp*CS were purified by the same method, except that cell extracts contained *Sc*CS were heated to 75 °C for 20 min[3], and the two proteins were eluted with 250 mM imidazole[4].

Recombinant *Tt*CS, *h*CS and *At*CS were also purified in this way while all buffer used contained 50 mM Tris-HCl (pH 8.0), 150 mM NaCl. The cell extracts of *Tt*CS was heated to 85 °C for 15 min[5], while *h*CS and *At*CS were not heated and were eluted with 250 mM imidazole.

**Crystallization**

Crystallization of purified *Da*CS, *Tt*CS and *h*CS was performed by the sitting-drop, vapor-diffusion method. Each drop contained 1 μL protein solution of which concentration was 10 mg/mL and 1 μL reservoir solution (reagents from Hampton Research).

The crystal of *Da*CS-citrate complex was grown in reservoir solution containing 0.2 M ammonium citrate tribasic (pH 7.0), 20% w/v Polyethylene glycol 3,350 at 16 °C for 5 days, while the crystal of *Da*CS-intermediate complex in 0.1M HEPES (pH 7.5) and 40% v/v Polyethylene glycol 400 for 4 days, and the *Da*CS-oxaloacetate complex crystal in 0.2 M Potassium chloride, 0.1 M HEPES (pH 7.5) and 35% v/v Pentaerythritol propoxylate (5/4 PO/OH) for 4 days .

The crystals of *Tt*CS-complexs were obtained by co-crystallizing with 2 mM CoA and 2 mM citric acids (Sigma). CoA-binding *Tt*CS crystals were growing in reservoir solution containing 0.1 M BIS-TRIS (pH 6.5), 2.2 M (*Tt*CS-citrate/CoA complex) or 2.4 M Ammonium sulfate (*Tt*CS-intermediate complex) at 16 °C for 3 days, while acetyl-CoA binding *Tt*CS crystals (*Tt*CS-oxaloacetate/acetyl-CoA complex) growing in 0.1 M BIS-TRIS (pH 6.5), 2.4 M Ammonium sulfate at 60 °C for 7 days.

The crystals of hCS were grown in reservoir solution containing 0.2 M $CH_3COONH_4$ (pH=7.1), 20% w/v PEG 3,350 refer to Schlachter, C. R. *et al*[6]. Crystals grew at 16 °C for 2 months or longer.

**Data collection, structure determination and refinement**

Crystal of *Da*CS-citrate was soaked in reservoir solution containing 20% glycerol.

Crystal of *Da*CS-intermediate complex was soaked in reservoir solution containing 20% glycerol, 50 mM citric acid and 50 mM CoA. Crystals of *Da*CS-oxaloacetate complex were soaked in reservoir solution containing 16.7 mM citric acid and 16.7 mM coenzyme A, then soaked in reservoir solution containing citric acid and CoA of the same concentrations and 20% glycerol.

The crystal of *Tt*CS-intermediate complex was soaked in reservoir solution containing 50 mM citrate (pH 5.0) and 6.5 mM CoA before soaked in the same solution with an additional 20% glycerol, while the crystal of *Tt*CS-citrate/CoA complex only soaked in the latter solution. The crystal of *Tt*CS-oxaloacetate/acetyl-CoA complex was soaked in reservoir solution containing 10 mM citrate (pH 5.0) and 13 mM CoA, then soaked in same solution containing 20% glycerol in addition.

For *h*CS-substrates or intermediates complexes, *h*CS crystals were soaked in reservoir solution containing 30 mM oxaloacetate and 50 mM acetyl-CoA, then soaked in reservoir solution containing same concentration of oxaloacetate and acetyl-CoA, and 20% glycerol. For *h*CS-oxaloacetate complex, crystal was soaked in reservoir solution containing 30 mM oxaloacetate and 30 mM acetyl-CoA, then soaked in same buffer containing 20% glycerol. For *h*CS-citrate complex, crystal was soaked in reservoir solution containing 50 mM citrate pH=5.0, then soaked in same buffer containing 20% glycerol.

All crystals were cryo-cooled in liquid nitrogen subsequently.

X-ray diffraction data sets for all the crystals were collected at the wavelength of 0.979 Å at BL10U2 (with Eiger-X 16M detector)[7] and BL18U1 (with Eiger-X 4M detector) in Shanghai Synchrotron Radiation Facility (SSRF) and then processed with the HKL3000 package[8] or XDS package[9].

The structures of *Da*CS and *Tt*CS were solved by molecular replacement using Phaser MR[10] with models predicted by alphafold2[11-12], while the structures of *h*CS-complex were solved by molecular replacement with apo-*h*CS structure (PDB: 5UZR) and refined by Refmac5[13] in CCP4 suit[14]. And their molecular models were built using COOT[15].

Cell parameters of *h*CS-OAA, *h*CS/OAA/AcCoA and *h*CS-product structures affected by conformational changes from the substrates binding leads to the difference in R factors.

All the figures of structures were prepared using PyMol (https://pymol.org), while figures involving chemical structures were prepared by ChemDraw (https://www.chemdraw.com.cn/).

**Activity assay of coenzyme A production**

The forward initial velocity of citrate synthases was measured by quantifying the thiol group of produced coenzyme A with 5,5'-dithiobis-(2-nitrobenzoic acid) (DTNB) spectrophotometrically at 412 nm[16] referring to Takuro, N. et al.[17]

The 100 µL reaction mixture mixed on ice contained 3 mM oxaloacetate acid, 0.5 mM acetyl-CoA (Sigma), 100 mM HEPES-NaOH (pH 7.5) and 0.05 µg citrate synthase (or not) and was incubated at 25 °C, 37 °C, 50 °C, 70 °C or 90 °C for 2 minutes. Then proteins were removed with Amicon Ultra-0.5mL 30kDa centrifugal filters (Milipole) at 4 °C to stop reactions.

20 μL filtrate of each group was added into 100 μL 1 mM DTNB which had been diluted with 1/15 M $Na_2HPO_4$-$KH_2PO_4$ (pH 7.27) from stock containing 0.086 M Tris, 0.09 M glycine and 4 mM EDTA (pH 8.0). After 20 minutes incubation at room temperature, $A_{412\ nm}$ of mixtures was measured. Simultaneously, a standard curve was determined with a concentration gradient of coenzyme A.

**UPLC measurements**

Acetyl-CoA produced in reverse reaction of citrate synthases were detected by ultra performance liquid chromatography (UPLC) according to Takuro, N. et al.[17]

The 200 μL reaction mixture contained 10 mM citric acid, 1 mM CoA, 100 mM Tris-HCl (pH 8.0 at 25 °C or at 70 °C) and 5 μg citrate synthase or not. After mixed on ice, the groups containing Tris-HCl (pH 8.0) were incubated at 25 °C or 37 °C for 2 hours, while the others at 50 °C, 70 °C or 90 °C. For detecting the effect of $H_2O_2$, the 200 μL mixture contained 1 μM $H_2O_2$, citric acid and CoA of the same concentrations, 100 mM Tris-HCl (pH=8.0 at 70 °C) and 100 μg *Da*CS (or not) and were incubated at 70 °C. Then reactions were stopped on ice and proteins were removed with Amicon Ultra-0.5mL 30kD centrifugal filters (Milipole) at 4 °C.

After centrifuged at 4 °C, 16,200 *g* for 10 minutes, 10 μL supernatant of each sample was loaded on a reversed-phase C18 column (Waters, ACQUITY UPLC ®HSST3, 1.8 μm, 2.1 x 100 mm column) in the Acquity UPLC H-Class system (Waters). CoA compounds inside were eluted with a methanol-20 mM $NaH_2PO_4$ (pH 5.3) gradient at a flow rate of 0.3 mL/min and detected at 254 nm. During elution, the volume fraction of methanol linearly changed from 10% to 15% in 2 minutes, then from 15% to 20% in

5 minutes, finally from 20% to 22% in 1 minute. Then the column was re-equilibrated with 10% methanol in 20 mM $NaH_2PO_4$ (pH 5.3) at the same flow rate for 10 minutes.

**NMR measurements**

Nuclear magnetic resonance (NMR) was used to detect the production of acetyl-CoA by *Da*CS.

The 600 μL reaction mixture contained 10 mM citric acid-2, 4-$^{13}C_2$ (Sigma), 10 mM CoA, 100 mM $Na_2HPO_4$-$KH_2PO_4$ (pH 7.0) and 300 μg citrate synthase (or not). For detecting the effect of free radical scavengers, 1 mM *L*-Ascorbic acid (Sigma) or 1 mM melatonin were added. After mixed on ice, all groups were incubated at 70 °C or 37 °C for 2 hours. Then proteins were removed with Amicon Ultra-0.5mL 30kD centrifugal filters (Milipole) at 4 °C.

400 μL filtrate each was added into 5 mm NMR sample tube mixing with 20 μL $D_2O$, while the 150 μL acetyl-CoA standard sample was added into 3 mm NMR sample tube with 20 μL $D_2O$.

$^{13}C$ NMR spectra were recorded on an Avance III 600MHz spectrometer (Bruker) at room temperature, each sample was scanned 258 times. Data were analyzed and figures were made using Topspin (Bruker).

**EPR measurements**

The possible free radicals in the reverse reaction catalyzed by *Da*CS were detected using electron paramagnetic resonance (EPR).

The 20 μL reaction mixture contained 10 mM citric acid, 10 mM CoA, 10 μM *Da*CS (or not) in 100 mM $Na_2HPO_4$-$KH_2PO_4$ (pH 7.0), or 10 μM citric acid, 10 μM CoA and

100 μM *Da*CS (or not) in same buffer and were mixed on ice. Then 2 μL 1 M 5, 5-Dimethyl-1-pyrroline N-oxide (DMPO) was added. All mixtures were incubated at 70 °C for 10 minutes before cooled and transferred into quartz capillary tubes.

EPR spectra were measured on a Bruker EMX plus/10-12 spectrometer (Bruker BioSpin GmbH) at room temperature with the following settings. Magnetic field scanning ranged from 3260 to 3460 Gauss with 1000 points resolution. The frequency of microwave was 9.438 GHz at 2.19 mW power. For receiver, modulation frequency was 100.00 kHz, and modulation amplitude was 1.00 Gauss. Each reaction mixture was scanned 10 times and the sweep time of each scan was 20.00 seconds.

**Raman Spectroscopy.**

The Raman spectra were detected at 785 nm using laser confocal Raman microscope (LabRAM HR Evolution) at room temperature (298K), and were collected from *h*CS crystals soaked in 2 μL droplet containing 30 mM oxaloacetate and 50 mM acetyl-CoA in reservoir solution. Data were collected after droplet containing substrates was added on crystals, using a laser power of 17 mW and exposure times of 100 sec. Control group data were collected under the same conditions, with unrelated crystal soaked in droplet containing 30 mM oxaloacetate and 50 mM acetyl-CoA (substrates) or 30 mM citrate and 50 mM CoA (products), or *h*CS crystals soaked in reservoir solution. Baseline correction were carried out using Origin (http://www.originlab.com).

The two bands at 745 $cm^{-1}$ and 749 $cm^{-1}$ are single-point peak observed in spectra of different crystals at similar reaction times, and both their signal intensity is close to that of surrounding peaks.

### Intermediates structures of *h*CS

The series structures of intermediates show the reaction intermediate states and the generation and break of chemical bonds in citrate synthesis process. The binding site of oxaloacetate/citrate consists of $His^{265}$, $His^{301}$, $His^{347}$, $Arg^{356}$, $Arg^{428}$ and $Arg^{448*}$ (* means from another subunit) with $Asp^{402}$ participating in catalysis located nearby, while the acetyl-CoA/CoA binds with loop 338-355 and loop 385-401.

The structure of *h*CS-oxaloacetate complex (PDB: 8ZVL) shows the free conformation of oxaloacetate (Supplementary Fig. 14a). The two carboxyl group of oxaloacetate bind with $Arg^{356}$ or $His^{265}$ and $Arg^{448*}$, and the C3-ketone group (C3-O3, Supplementary Fig. 14a) points towards $His^{301}$, showing the state before the reaction.

The structure of *h*CS complexed with oxaloacetate and acetyl-coenzyme A (PDB: 8ZW1) represents the initial state at the start of the reaction. The conformation of oxaloacetate allows reaction to start (Supplementary Fig. 14b), in which the C3-ketone group (C3-O3, Supplementary Fig. 14b) points to the acetyl-group of acetyl-CoA (Supplementary Fig. 15d-f).

In structure of *h*CS-intermediate 1 (PDB: 8ZVM), the electron density shows acetyl analogue, oxaloacetate and CoA binding in active site. The oxygen of acetyl analogue ($O1^{(ACE)}$, Supplementary Fig. 14c) has interaction with ketone of oxaloacetate ($O3^{(OAA)}$, Supplementary Fig. 14c) with 2.2 Å distance between the two oxygen (Supplementary Fig. 14c, Supplementary Fig. 16a-c). The water molecular nearby $Asp^{402}$ contributes to catalyzing the hydrolysis of thioester bond in acetyl-CoA (Supplementary Fig. 14c, Supplementary Fig. 16a-c) and it is reasonable that this acetyl analogue is produced by the hydrolysis of acetyl-CoA. The binding of oxaloacetate and CoA is similar to that in *h*CS/oxaloacetate/acetyl-CoA complex.

In *h*CS-intermediate 2 structure (PDB: 8ZVR), the electron density observed near CoA can be matched as acetyl group and oxaloacetate rather than citrate (Supplementary Fig. 16d-f, Supplementary Fig. 18). The distance between acetyl oxygen ($O1^{(ACE)}$, Supplementary Fig. 14d) and ketone oxygen of oxaloacetate ($O3^{(OAA)}$, Supplementary Fig. 14d) is 1.4 Å, and the distance between $-CH_3$ of acetyl group ($C2^{(ACE)}$, Supplementary Fig. 14d) and ketone carbon of oxaloacetate ($C3^{(OAA)}$, Supplementary Fig. 14d) is 2.5 Å (Supplementary Fig. 14d). The interaction between the two oxygen atoms is suspected as peroxide bond, and it also indicates that C-C bond has not been formed between the two carbon atoms (Supplementary Fig. 16d-f). The water molecular, or hydroxyl group, which is located nearby $Asp^{402}$ and acetyl analogue (Supplementary Fig. 14d), is possibly involved in catalysis.

In structure of *h*CS-intermediate 3 (PDB: 8ZVT), the electron density shows acetate analogue and oxaloacetate in active site (Supplementary Fig. 16g-i, Supplementary Fig. 19). The 1.9 Å distance between carboxyl oxygen of acetate analogue ($O1^{(ACY)}$, Supplementary Fig. 14e) and O3 of oxaloacetate indicates the formation of O-O bond which precedes the generation of C-C bond, while the distance between methyl carbon ($C2^{(ACY)}$, Supplementary Fig. 14e) and C3 of oxaloacetate is 2.1 Å (Supplementary Fig. 14e).

In structure of *h*CS-intermediate 4 (PDB: 8ZVU), the electron density shows citrate analogue located near CoA in the active site (Supplementary Fig. 14f, Supplementary

Fig. 16j-l). Its O3, C3, C5, C6 and O6 are coplanar and suspected to have formed five-membered ring. The electron density (Supplementary Fig. 16j-l) and the 2.0 Å distance between O3 and O6 (Supplementary Fig. 14f) indicates that a possible peroxide bond exists between the two oxygen atoms.

All *h*CS-intermediate structures are obtained by soaking apo-crystals in oxaloacetate and acetyl-CoA without citrate or other ligands. These ligands observed in those structures represent intermediates in citrate synthesis reaction.

In structure of *h*CS-byproduct complex (PDB: 8ZVW), hydroxyl is observed near oxaloacetate in the active site without acetyl analogue or acetate analogue (Supplementary Fig. 14h). The result indicates that the synthesis of citrate has not been completed but the thioester bond in acetyl-CoA has been hydrolyzed. This structure possibly represents reaction-failed state.

The structure of *h*CS citrate complex (PDB: 8ZVV) was obtained by soaking *h*CS crystal into solution containing citrate. In this structure, the C6-carboxyl of citrate are not coplanar with O3, C3 and C5. All the atoms of citrate obey stereochemical constrains without forming plane like that in *h*CS-intermediate 4 and there is not strong interaction between O3 and O6 (Supplementary Fig. 17a-c) according to the 2.9 Å distance (Supplementary Fig. 14g). This structure represents the final state of reaction.

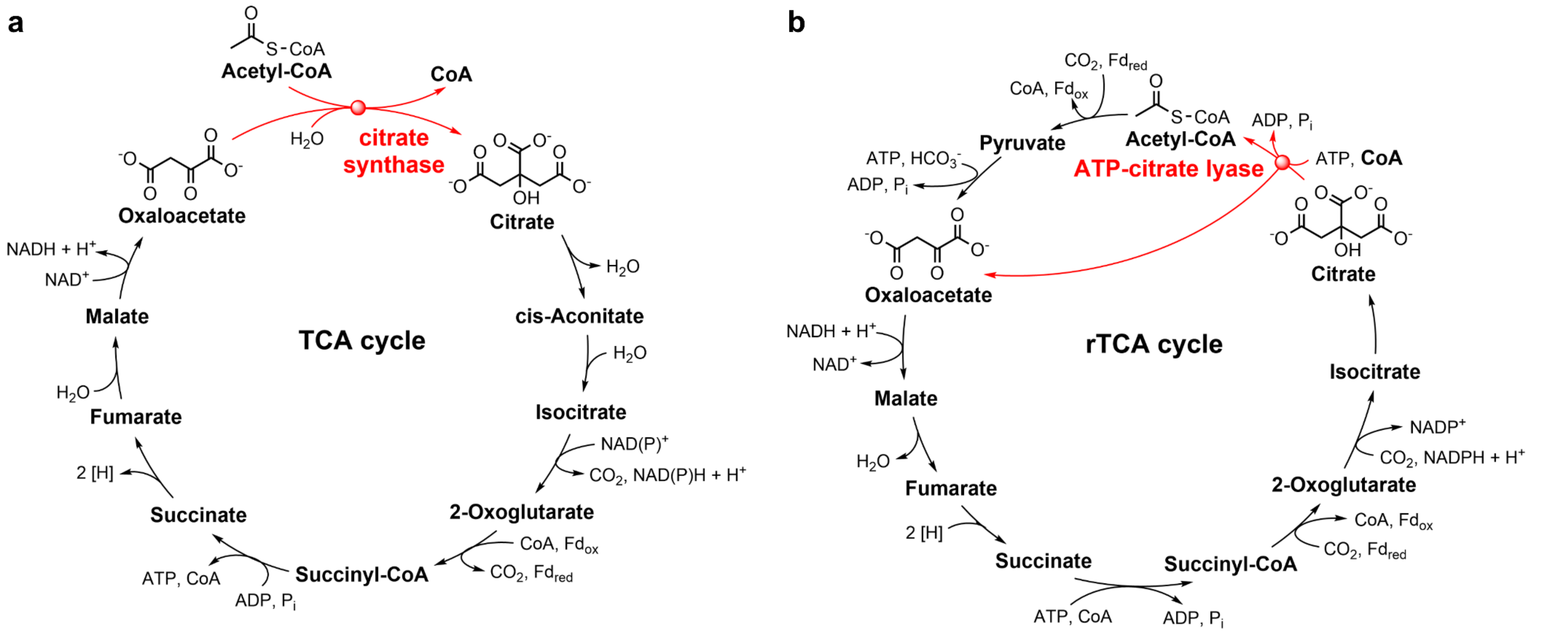


**Supplementary Fig. 1 | TCA cycle and reverse TCA cycle.**

**a**, General oxidative tricarboxylic acid (TCA) cycle, with citrate synthesis reaction shown in red. **b,** Reverse tricarboxylic acid (rTCA) cycle according to that in *Hydrogenobacter thermophiles*[18], with citrate cleavage reaction shown in red.

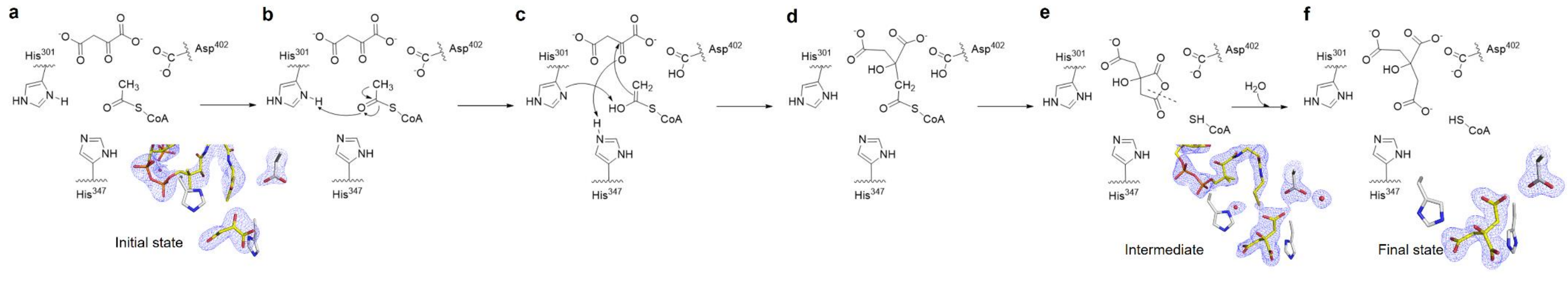


**Supplementary Fig. S2 | Classical catalytic mechanism of *h*CS.** Structures are shown nearby.

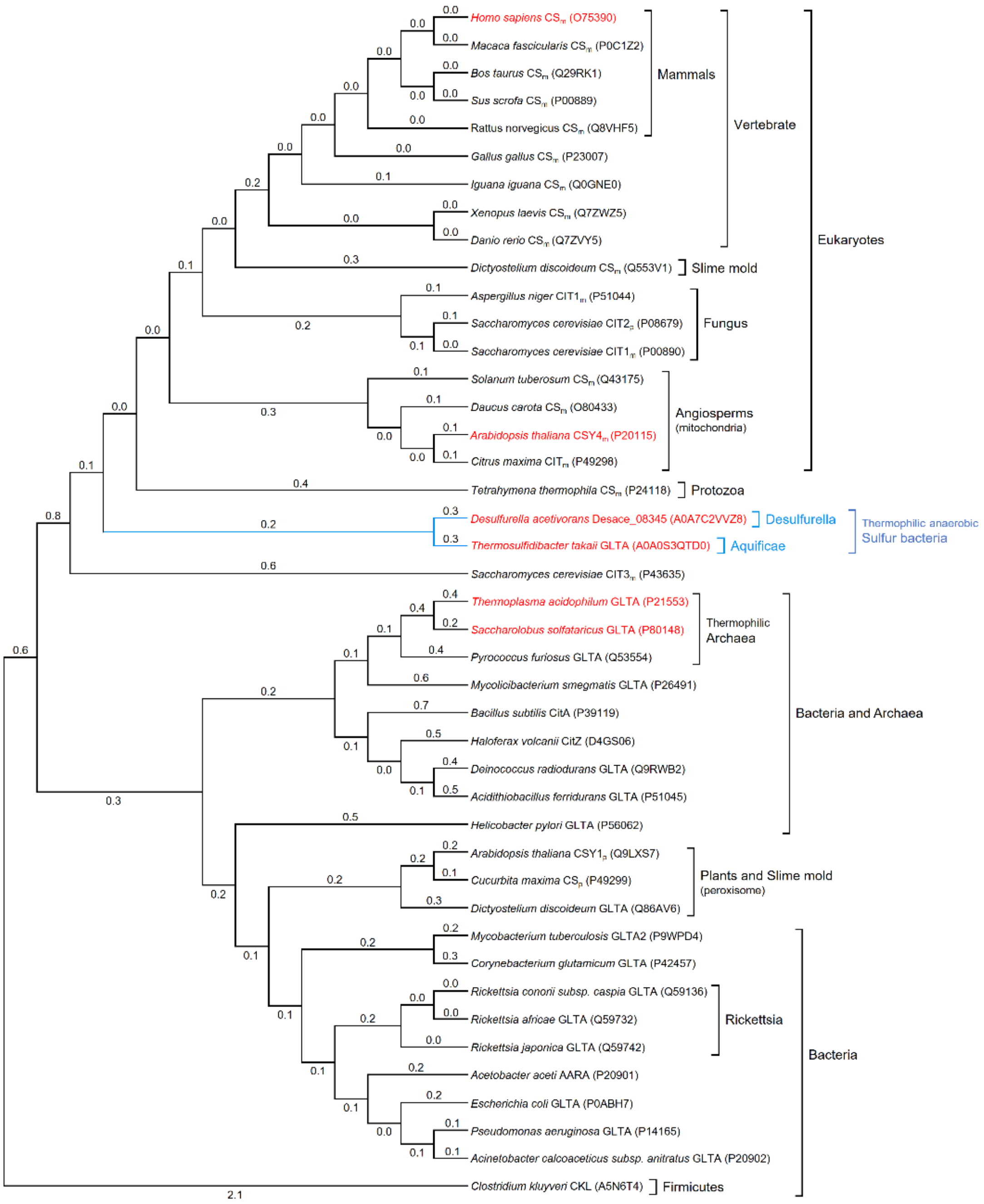


**Supplementary Fig. 3 | Phylogenetic tree of citrate synthases.**

43 amino acid sequences of citrate synthases were aligned using Clustal Omega[19-21] on EBI web server[22] (www.ebi.ac.uk/Tools/msa/clustalo), and the evolutionary analysis was conducted in MEGA7[23]. The evolutionary history was inferred using Maximum Likelihood method based on JTT matrix-based model[24]. The log likelihood of the unrooted tree is (-14161.17). There was a total of 273 positions in the final dataset while

all positions containing gaps and missing data were eliminated. Each citrate synthase involved in phylogenetic tree was shown with its organism source, gene name and the Uniprot ID given in parentheses. For eukaryotic citrate synthase, its subcellular location was annotated with a letter "m" standing for mitochondria or a "p" for peroxisome as subscript. The six citrate synthases involved in activity assays were marked in red, while the two roTCA citrate synthases among them were annotated with blue lines.

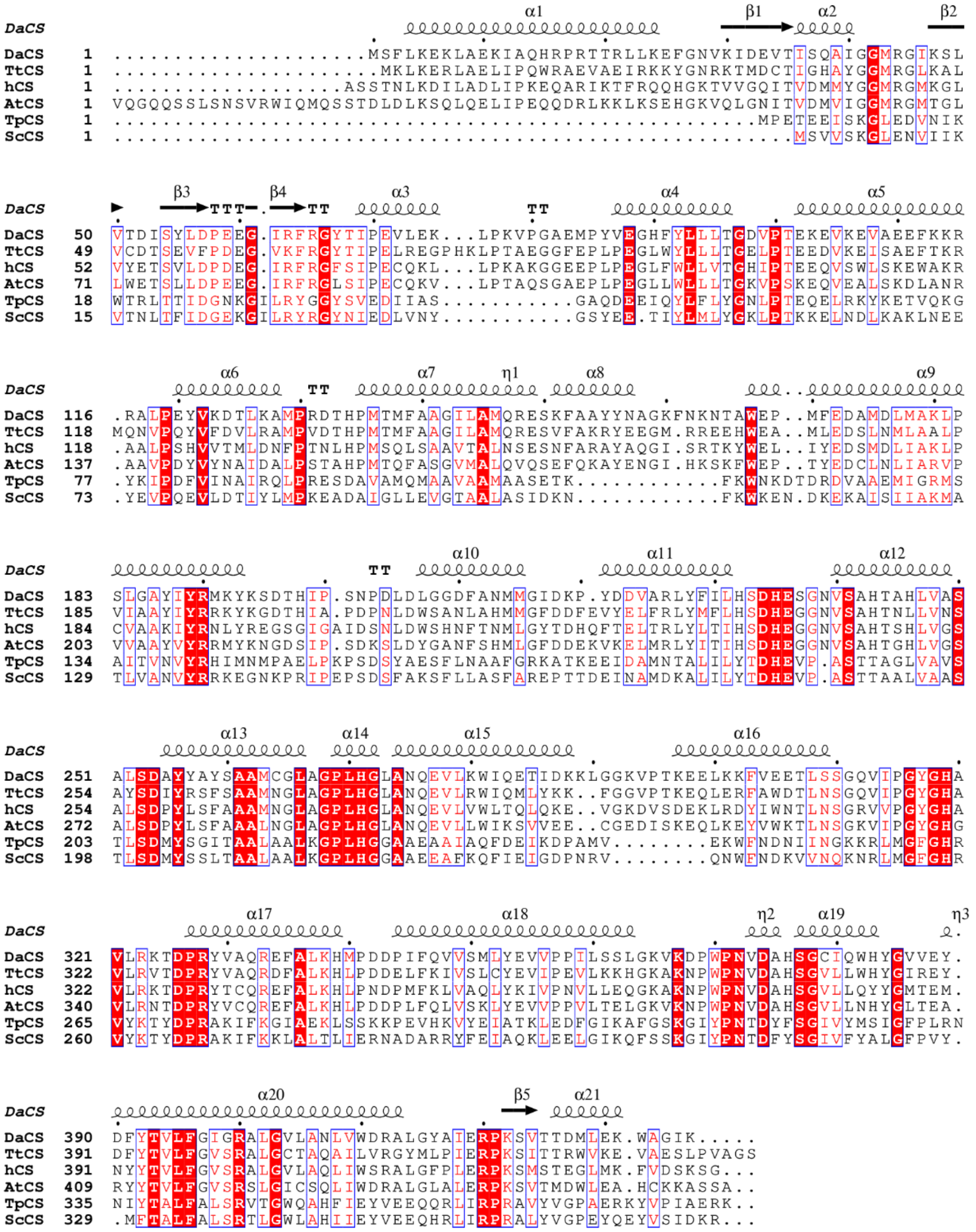

**Supplementary Fig. 4 | Sequence alignment of *Da*CS, *Tt*CS, *h*CS, *At*CS, *Tp*CS and *Sc*CS.**

The amino acid sequences of six citrate synthases without signal peptide were aligned using Clustal Omega[19-21] on EBI web server[22] (www.ebi.ac.uk/Tools/msa/clustalo), and the result were visualized by Espript 3.0[25] (espript.ibcp.fr/ESPript/cgi-bin/ESPript.cgi).

Each sequence’s phylogenetic position are shown in Supplementary Fig. 3. The strictly identical residues are in white on a red background, while equivalent residues whose global score is higher than 70% are colored in red and framed in blue. The secondary structure of *Da*CS, which is predicted by its tertiary structure, was shown above the sequences, with a letter “α” standing for α–helix, “β” for β–strand, “η” for $3_{10}$-helix and “TT” for turn.

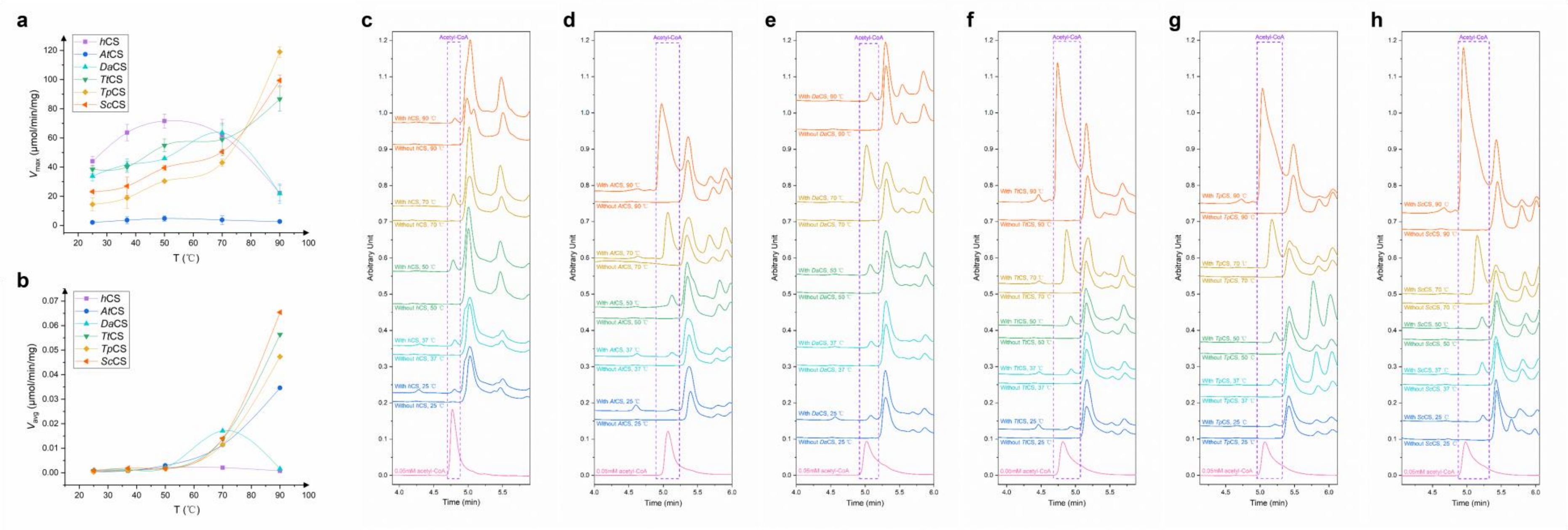


**Supplementary Fig. 5 | Forward and reverse reaction velocity and acetyl-coenzyme A production of citrate synthases at different temperatures.**

**a**, Maximum forward reaction velocity for *h*CS, *At*CS, *Da*CS, *Tt*CS, *Tp*CS and *Sc*CS at 25 °C, 37 °C, 50 °C, 70 °C or 90 °C. Data are plotted as mean ± standard error of the mean generated from n = 3 independent biological replicates. **b**, Average reverse reaction velocity in two hours for *h*CS, *At*CS, *Da*CS, *Tt*CS, *Tp*CS and *Sc*CS at 25 °C, 37 °C, 50 °C, 70 °C or 90 °C, according to the peak area of acetyl-CoA in UPLC spectra. **c~h**,

UPLC analysis of acetyl-CoA production at 254 nm catalyzed by *h*CS **c**, *At*CS **d**, *Da*CS **e**, *Tt*CS **f**, *Tp*CS **g** and *Sc*CS **h** at 25 ℃, 37 ℃, 50 ℃, 70 ℃ or 90 ℃. 0.05 mM acetyl-CoA was used as standard, and reaction mixtures without enzyme are shown as negative control in each schematic. Spectra are moved along horizontal axis within 0.02 unites to align the peaks of acetyl-CoA.

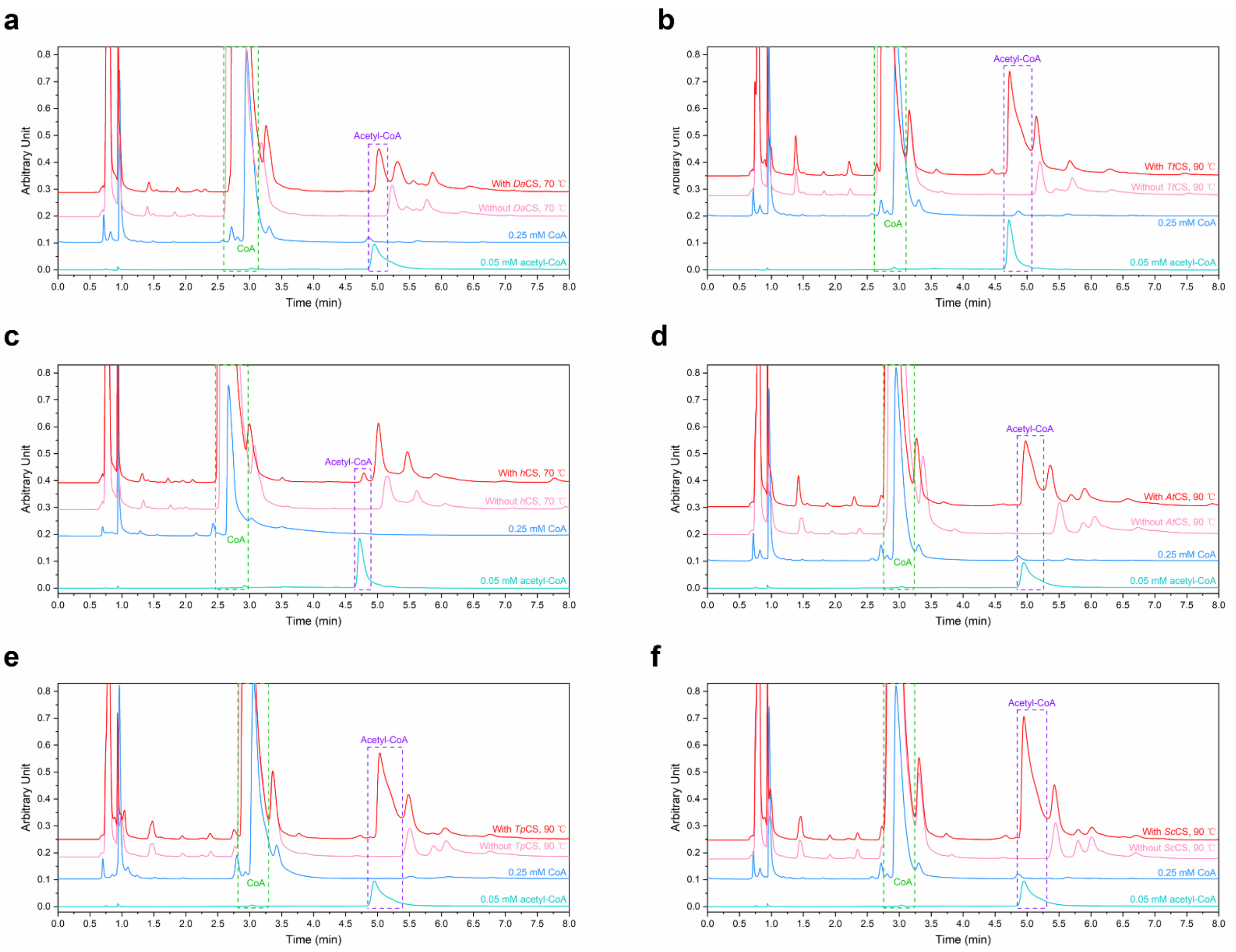
a
b
c
d
e
f
Arbitrary Unit
Time (min)
CoA
Acetyl-CoA
With DaCS, 70 ℃
Without DaCS, 70 ℃
0.25 mM CoA
0.05 mM acetyl-CoA
With TtCS, 90 ℃
Without TtCS, 90 ℃
With hCS, 70 ℃
Without hCS, 70 ℃
With AtCS, 90 ℃
Without AtCS, 90 ℃
With TpCS, 90 ℃
Without TpCS, 90 ℃
With ScCS, 90 ℃
Without ScCS, 90 ℃

**Supplementary Fig. 6 | UPLC spectra of reverse reaction mixture for citrate synthases.**

**a**, UPLC analysis of CoA compounds at 254 nm in reverse reaction mixtures of *Da*CS at 70 ℃. **b**, *Tt*CS at 90 ℃. **c**, *h*CS at 70 ℃. **d**, *At*CS at 90 ℃. **e**, *Tp*CS at 90 ℃. **f**, *Sc*CS at 90 ℃. For each citrate synthases, the spectrum with highest acetyl-CoA production was selected. 0.25 mM CoA and 0.05 mM acetyl-CoA were used as standard, and the reaction mixture without enzyme is shown as negative control in each schematic.

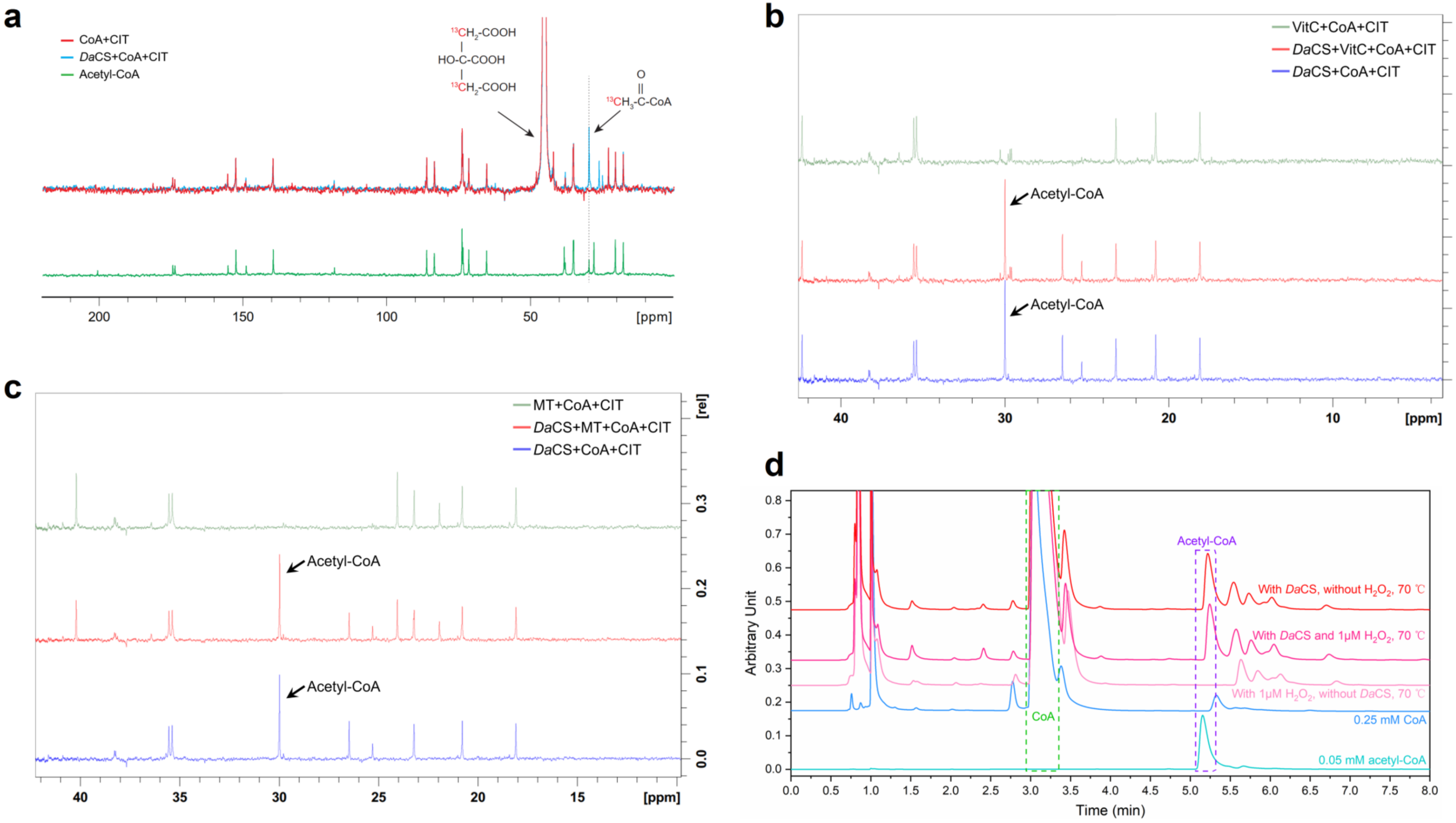
a
CoA+CIT
DaCS+CoA+CIT
Acetyl-CoA
13CH2-COOH
HO-C-COOH
13CH2-COOH
O
13CH3-C-CoA
200
150
100
50
[ppm]
b
VitC+CoA+CIT
DaCS+VitC+CoA+CIT
DaCS+CoA+CIT
Acetyl-CoA
Acetyl-CoA
40
30
20
10
[ppm]
[rel]
c
MT+CoA+CIT
DaCS+MT+CoA+CIT
DaCS+CoA+CIT
Acetyl-CoA
Acetyl-CoA
40
35
30
25
20
15
[ppm]
[rel]
d
Arbitrary Unit
Time (min)
Acetyl-CoA
CoA
With DaCS, without H2O2, 70 ℃
With DaCS and 1μM H2O2, 70 ℃
With 1μM H2O2, without DaCS, 70 ℃
0.25 mM CoA
0.05 mM acetyl-CoA

**Supplementary Fig. 7 | The effect of free radical scavengers and hydroxyl radical provider on acetyl-coenzyme A production catalyzed by *Da*CS.**

**a**, Complete $^{13}C$ NMR spectrum of reverse reaction mixture containing *Da*CS, [2, 4-$^{13}C$] citric acid and unlabeled CoA. 30 ppm $^{13}C$-methyl peak of produced [1-$^{13}C$] acetyl-CoA was detected. 75 mM unlabled acetyl-CoA was used as standard, and reaction mixture without enzyme is shown as negative control. **b**, $^{13}C$ NMR analysis of acetyl-CoA production catalyzed by *Da*CS at 70 °C with 1 mM *L*-ascorbic acid (VitC) as free radical scavenger. Reaction mixture without VitC and mixture without enzyme are shown as control. The area of 30 ppm [1-$^{13}C$] acetyl-CoA peak in spectrum of reaction mixture was nearly the same with or without VitC. **c**, $^{13}C$ NMR analysis of acetyl-CoA production catalyzed by *Da*CS at 70 °C with 1 mM melatonin (MT) as free radical scavenger. Reaction mixture without melatonin and mixture without enzyme are shown as control. The area of 30 ppm [1-$^{13}C$] acetyl-CoA peak in reaction mixture spectrum was nearly unchanged after adding melatonin. **d**, UPLC analysis of acetyl-CoA production at 254 nm by *Da*CS at 70 °C with 1 μM $H_2O_2$ for providing excess hydroxyl radical. 0.25 mM CoA and 0.05 mM acetyl-CoA were used as standard, while reaction mixture without $H_2O_2$ and mixture without enzyme are shown as control. The peak area of acetyl-CoA in spectrum of reaction mixture did not change after adding $H_2O_2$.

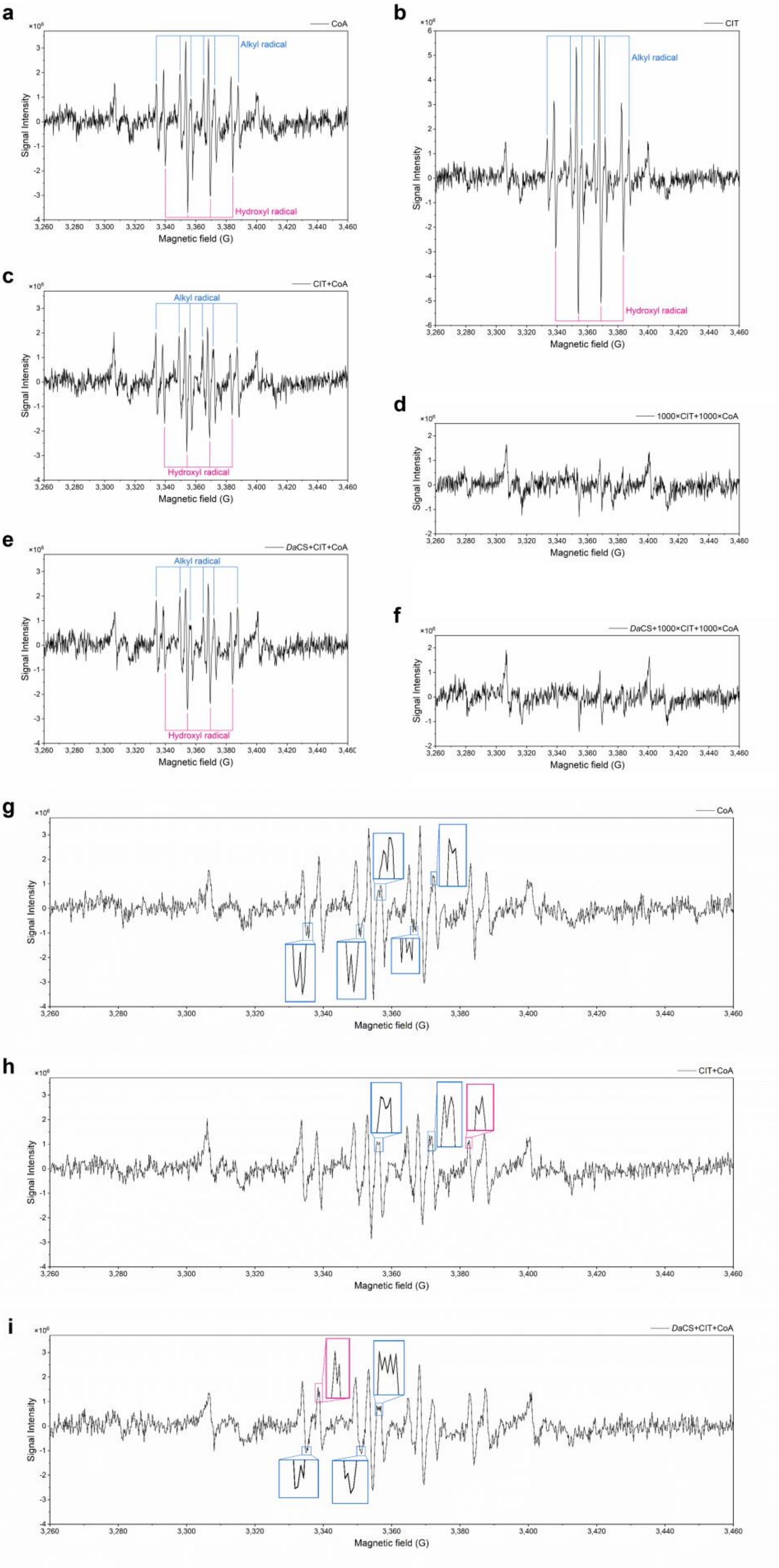
a
CoA
Alkyl radical
Hydroxyl radical
Signal Intensity
Magnetic field (G)
b
CIT
c
CIT+CoA
d
1000×CIT+1000×CoA
e
DaCS+CIT+CoA
f
DaCS+1000×CIT+1000×CoA
g
h
i

**Supplementary Fig. 8 | EPR characterization of free radical in heated reverse reaction system of *Da*CS.**

**a**, EPR spectrum of 10 μM CoA, incubated at 70 ℃ with 90.9 mM DMPO. Six-line signal from DMPO-alkyl radical adduct ($A_N$ = 15.63 G, $A_H^{\beta}$ = 22.66 G) are labeled in blue and four-line signal from hydroxyl radical ones ($A_N$ = 14.84 G, $A_H^{\beta}$ = 14.81 G) are annotated in pink. **b**, EPR spectrum of 10 μM citrate with labeled alkyl radical and hydroxyl radical signals. **c**, EPR spectrum of 10 μM CoA and 10 μM citrate with labeled radical species signals. **d**, EPR spectrum of 10 mM CoA and 10 mM citrate without characteristic signal, since radicals would be quenched with high concentration citrate. **e**, EPR spectrum of 100 μM *Da*CS,10 μM CoA and 10 μM citrate with labeled radical species signals. **f**, EPR spectrum of 100 μM *Da*CS, 10 mM CoA and 10 mM citrate without characteristic signal. **g**, Enlarged split peak tips in spectrum **a**. **h**, Enlarged split peak tips in spectrum **c**. **i**, Enlarged split peak tips in spectrum **e**. Tips from alkyl radical signal are shown in blue frame, while those from hydroxyl radical in pink. Values of g-factor and hyperfine splitting constants for spectrum **a**, **b**, **c** and **e** are shown in Supplementary Table 1.

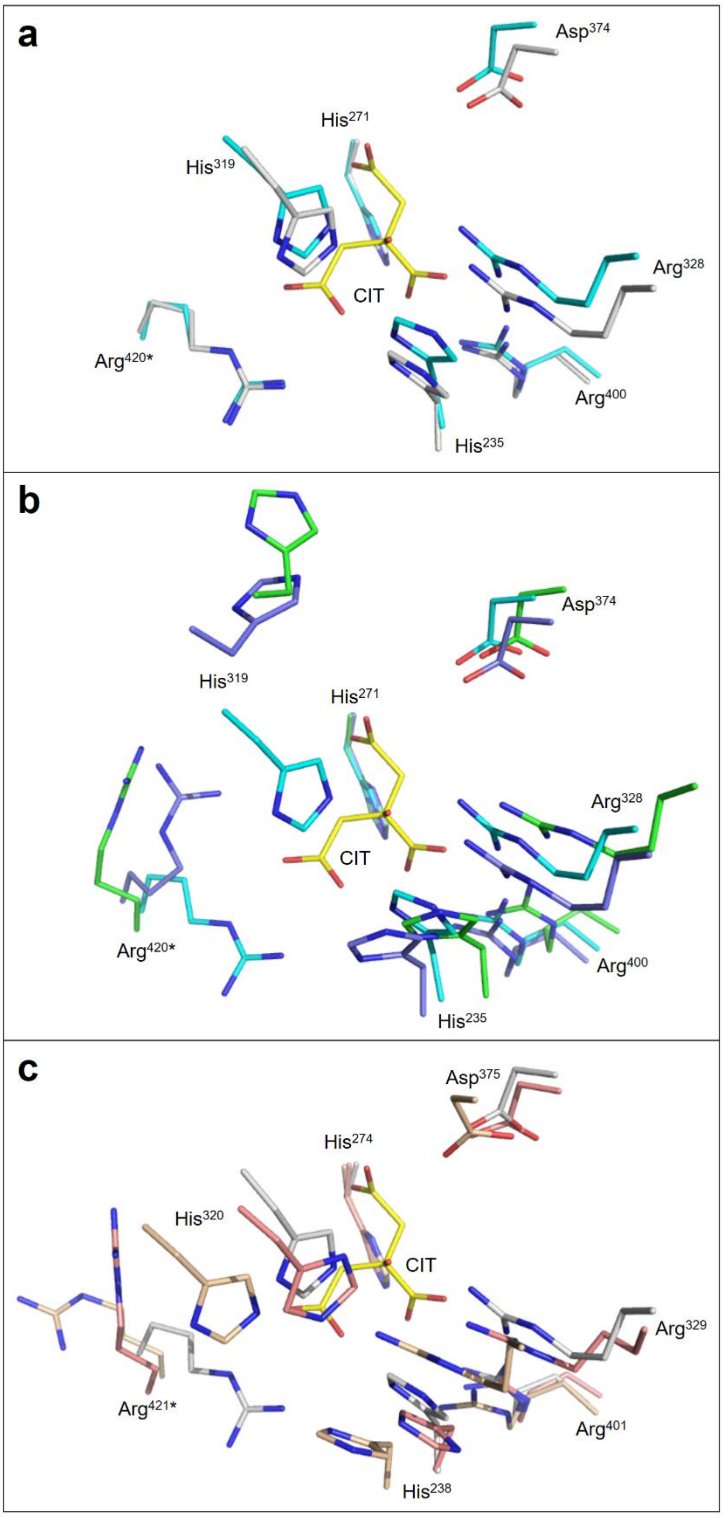


**Supplementary Fig. 9 | Active sites similarity among citrate synthases.**

**a**, Active site residues of *Da*CS (cyan, with label) (PDB: 8XU2) and *Tt*CS (gray) (PDB: 8XXD) with bound citrate (yellow) from *Da*CS structure. **b**, Active site residues of

*Da*CS (with label), *h*CS (slate) (PDB: 5UZR) and *At*CS (green) (PDB: 6K5V). **c**, Active site residues of *Tt*CS (with label), *Tp*CS (wheat) (PDB: 4YBO) and *Sc*CS (salmon) (PDB: 1O7X). All active sites from different citrate synthases are aligned against His$^{271}$ of *Da*CS while other conserved residues adopted similar position, except His$^{319}$ and Arg$^{420*}$ (number in *Da*CS) located in active loop and adopting relatively free conformation before citrate binding. Thus, their side chains swing into different orientations in *h*CS, *At*CS, *Tp*CS and *Sc*CS compared with *Da*CS and *Tt*CS.

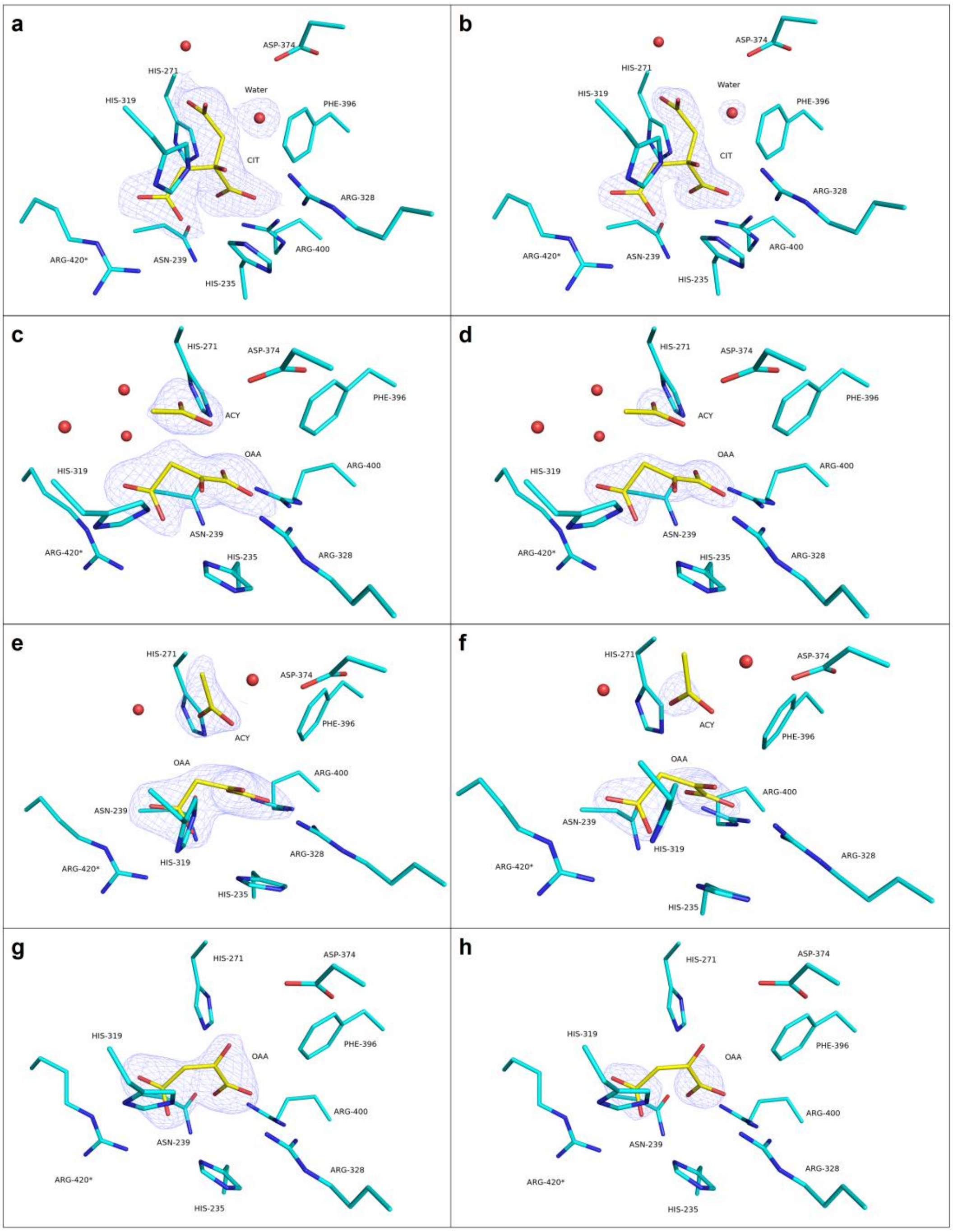


**Supplementary Fig. 10 | Structures of substrate, intermediate or product with electron density in active site of *Da*CS.**

**a**, Citrate (yellow) binding in active site of *Da*CS (cyan), with corresponding $2F_{obs}$ - $F_{calc}$ OMIT map (blue mesh), 1.0σ, from *Da*CS-citrate complex (PDB: 8XU2). **b**, Citrate binding same to **a**, with 2.0σ map. **c**, Oxaloacetate and acetate analogue in *Da*CS active site with 1.0σ map, from Chain E in *Da*CS-intermediates complex (PDB: 8XXC). **d**, Same to **c** with 2.0σ map. **e**, Oxaloacetate and acetate analogue of another conformation with 1.0σ map, in Chain F of *Da*CS-intermediates complex structure (PDB: 8XXC). **f**, Same to **e** with 2.0σ map. **g**, Oxaloacetate with 1.0σ map, from *Da*CS-oxaloacetate complex (PDB: 8XU1). h, Same to **g** with 2.0σ map.

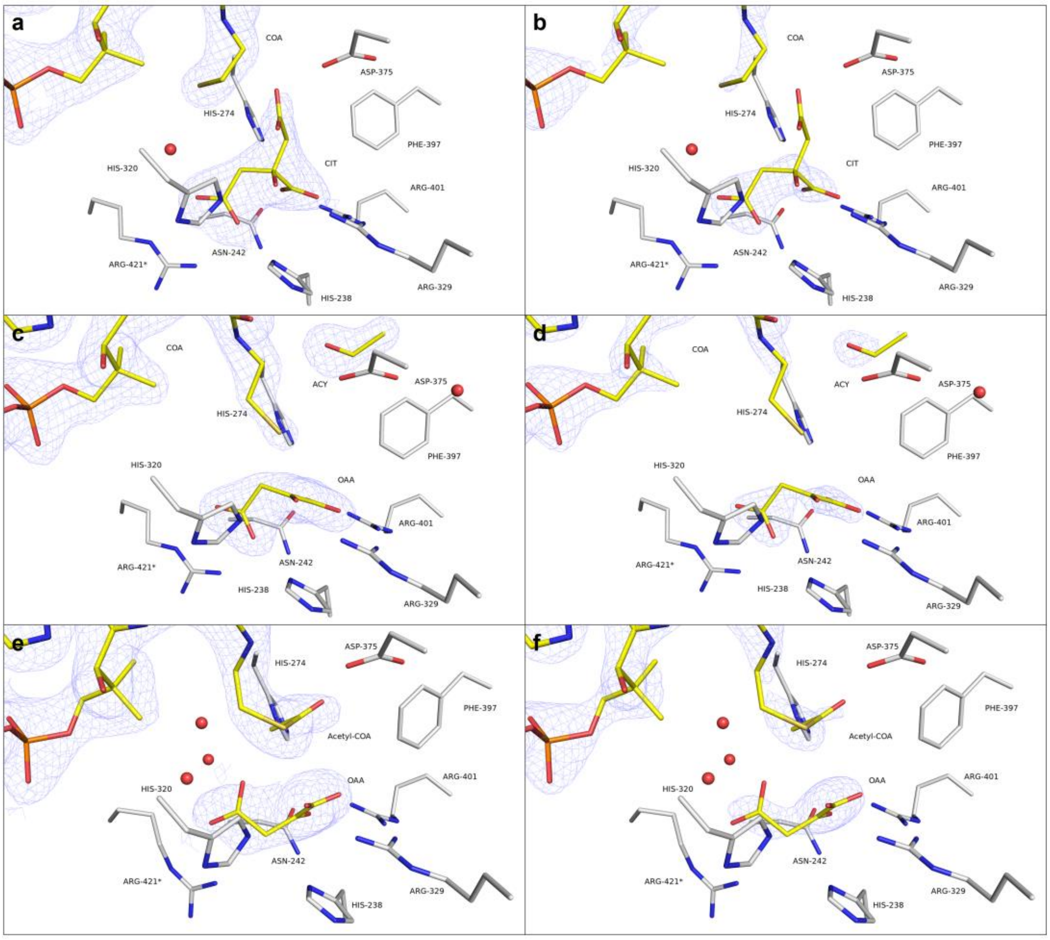


**Supplementary Fig. 11 | Structures of substrates, intermediates or products with electron density in active site of *Tt*CS.**

**a**, Citrate and coenzyme A (yellow) binding in active site of *Tt*CS (gray), with corresponding $2F_{obs}$ - $F_{calc}$ OMIT map (blue mesh), 1.0σ, from *Tt*CS-citrate/CoA complex structure (PDB: 8XXD). **b**, Same to **a**, with 2.0σ map. **c**, Oxaloacetate and acetyl group in active site with 1.0σ map, from Chain C in *Tt*CS-intermediates complex (PDB: 8XXE). **d**, Same to **c** with 2.0σ map. **e**, Oxaloacetate and acetyl-CoA with 1.0σ map, from Chain F in *TtCS*-oxaloacetate/acetyl-CoA complex (PDB: 8XXF). **f**, Same to **e** with 2.0σ map.

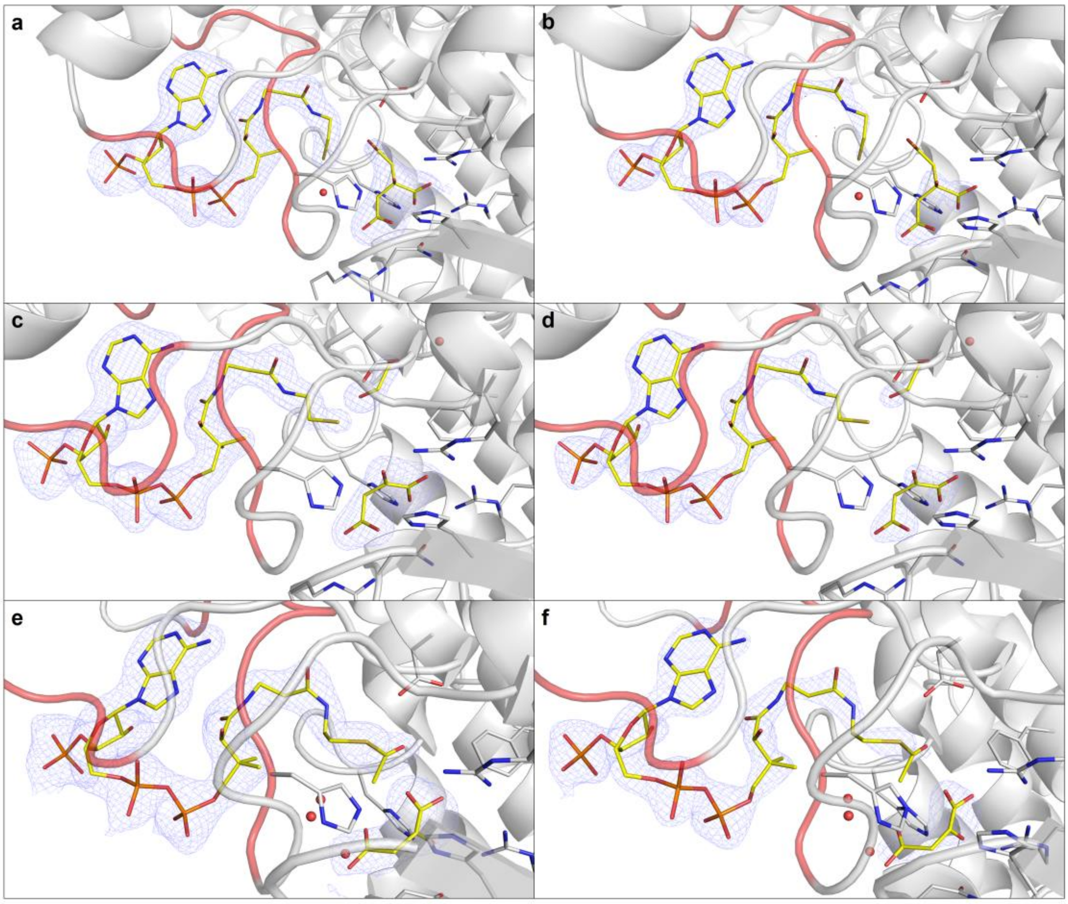


**Supplementary Fig. 12 | Complete coenzyme A compound structures with electron density binding with *Tt*CS.**

**a**, Coenzyme A and citrate (yellow) in CoA binding loops, loop 314-321 and loop 365-369 (red) of *Tt*CS (gray), with corresponding $2F_{obs} - F_{calc}$ OMIT map (blue mesh), 1.0σ, from *Tt*CS-citrate/CoA complex (PDB: 8XXD). **b**, Same to **a**, with 2.0σ map. **c**, CoA and oxaloacetate and acetyl group with 1.0σ map, from Chain C in *Tt*CS-intermediates complex structure (PDB: 8XXE). **d**, Same to **c** with 2.0σ map. **e**, Acetyl-CoA and oxaloacetate with 1.0σ map, from Chain F in *TtCS*-oxaloacetate/acetyl-CoA complex structure (PDB: 8XXF). **f**, Same to **e** with 2.0σ map.

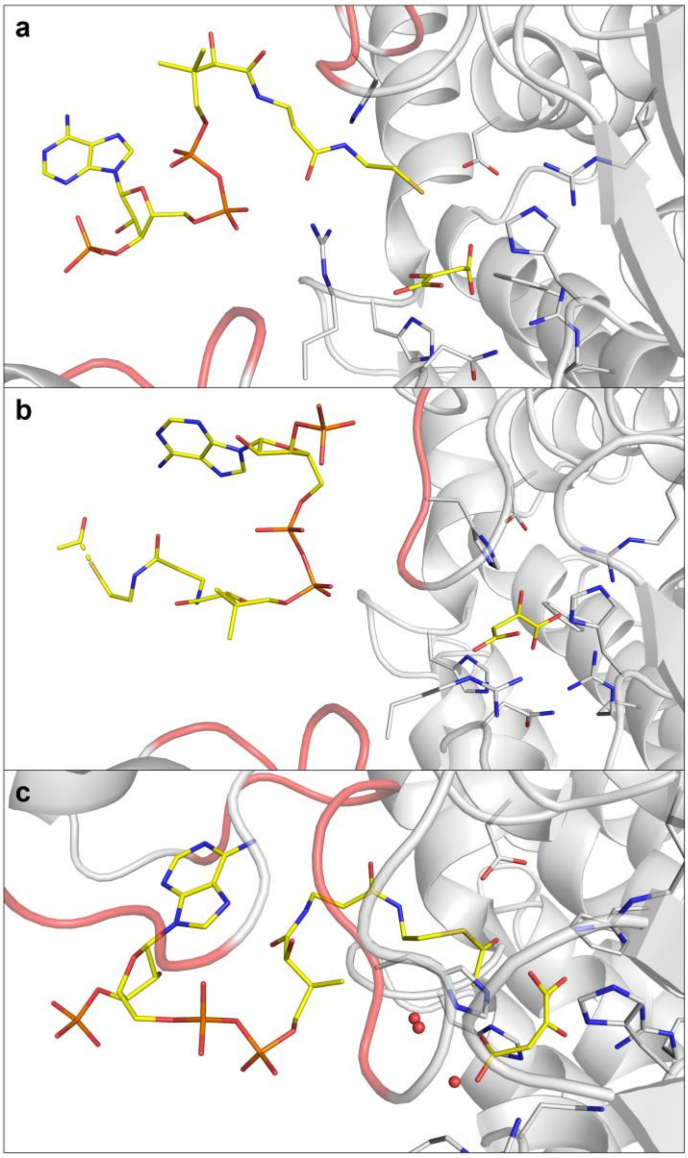


**Supplementary Fig. 13 | Different coenzyme A compound binding position in *Tt*CS structures.**

**a**, Coenzyme A (yellow) binding in loop 39-44 and loop 314-321 (red) of *Tt*CS (gray) with oxaloacetate (yellow) in active site, from Chain B of *Tt*CS-intermediate complex structure (PDB: 8XXE). **b**, Suspected acetyl-CoA binding in loop 39-44 and loop 314-321, from Chain A of *Tt*CS-oxaloacetate/acetyl-CoA structure (PDB: 8XXF). **c**, Acetyl-CoA binding in loop 314-321 and loop 365-369, from Chain F of *Tt*CS-oxaloacetate/acetyl-CoA structure (PDB: 8XXF).

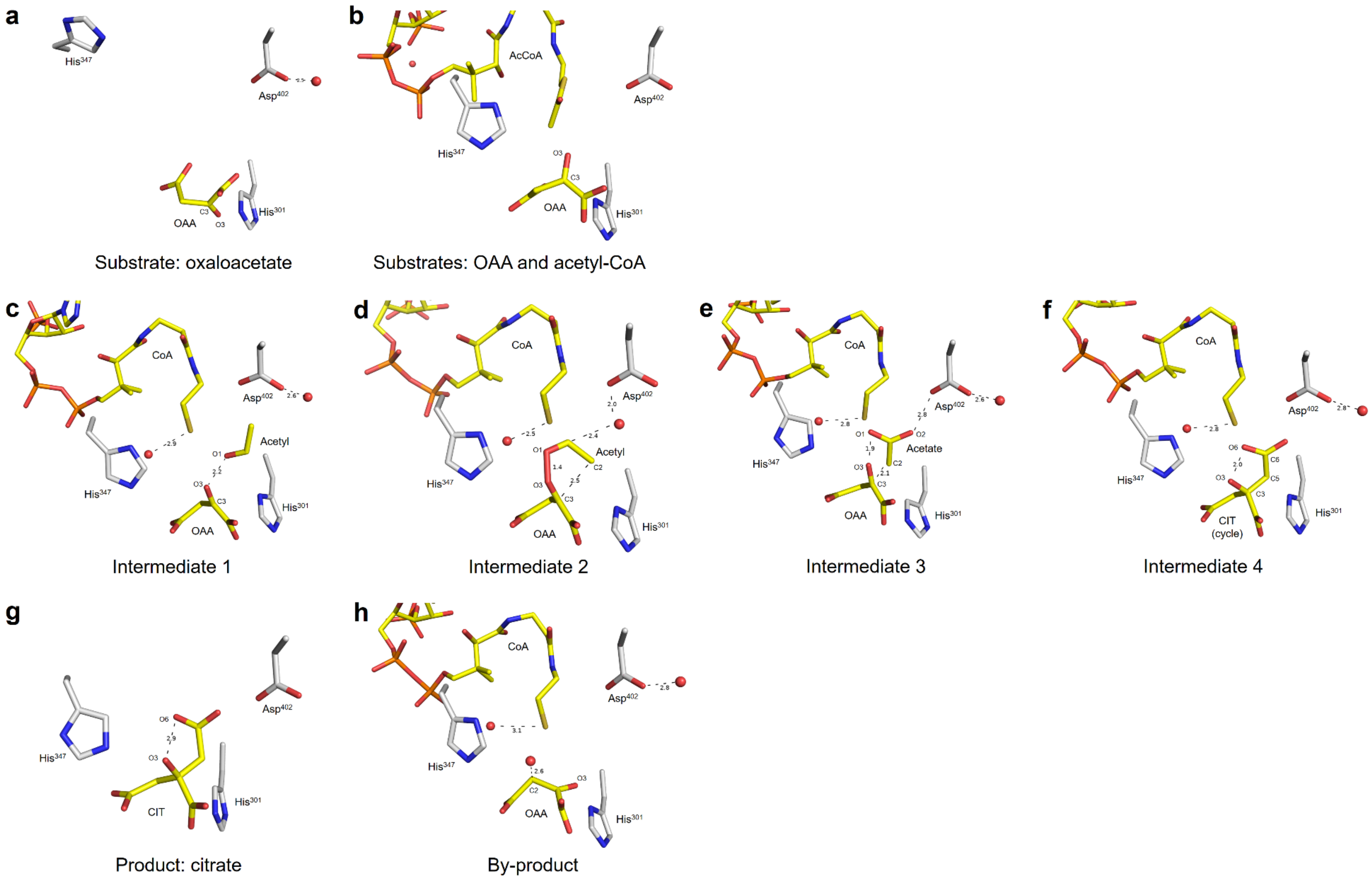
a
His$^{347}$
Asp$^{402}$
OAA
His$^{301}$
Substrate: oxaloacetate
b
AcCoA
Asp$^{402}$
His$^{347}$
OAA
His$^{301}$
Substrates: OAA and acetyl-CoA
c
CoA
Asp$^{402}$
Acetyl
His$^{347}$
OAA
His$^{301}$
Intermediate 1
d
CoA
Asp$^{402}$
Acetyl
His$^{347}$
OAA
His$^{301}$
Intermediate 2
e
CoA
Asp$^{402}$
Acetate
His$^{347}$
OAA
His$^{301}$
Intermediate 3
f
CoA
Asp$^{402}$
His$^{347}$
CIT
(cycle)
His$^{301}$
Intermediate 4
g
Asp$^{402}$
His$^{347}$
CIT
His$^{301}$
Product: citrate
h
CoA
Asp$^{402}$
His$^{347}$
OAA
His$^{301}$
By-product

**Supplementary Fig. 14 | Substrates, different intermediates, product and by-product found in *h*CS structure.** Ligand is colored in yellow with atoms labeled and distances annotated in dash. Proteins colored in gray. **a,** Active site of *h*CS-oxaloacetate complex structure (PDB: 8ZVL) **b,** *h*CS-substrates complex structure (PDB: 8ZW1). **c,** *h*CS-intermediate 1 structure (PDB: 8ZVM). **d,** *h*CS-intermediate 2 structure (PDB: 8ZVR). **e,** *h*CS-intermediate 3 structure (PDB: 8ZVT). **f,** *h*CS-intermediate 4 structure (PDB: 8ZVU). **g,** *h*CS-citrate structure (PDB: 8ZVV). **h,** *h*CS-byproduct structure (PDB: 8ZVW).

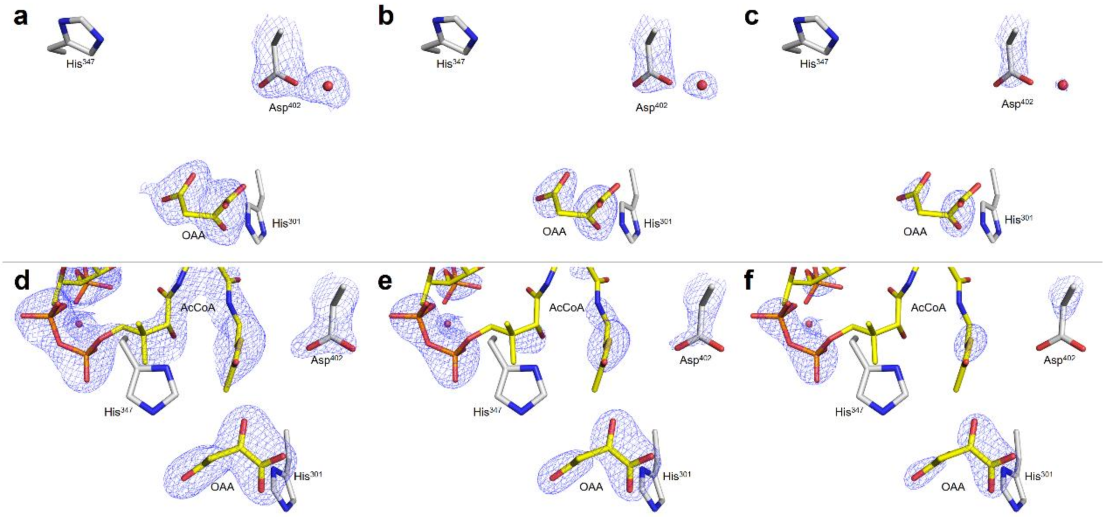


**Supplementary Fig. 15 | Structures of substrates in active site of *h*CS.**

**a,** Oxaloacetate (OAA, yellow) binding in *h*CS active site (gray) (PDB: 8ZVL). 2Fo-Fc omit map are shown as blue mesh in 1.0σ. **b,** Same to **a** with 1.5σ map. **c,** Same to **a** with 2.0σ map. **d,** Oxaloacetate and acetyl-CoA (AcCoA) as substrates in *h*CS active site (PDB: 8ZW1) with 1.0σ map. **e,** Same to **d** with 1.5σ map. **f,** Same to **d** with 2.0σ map.

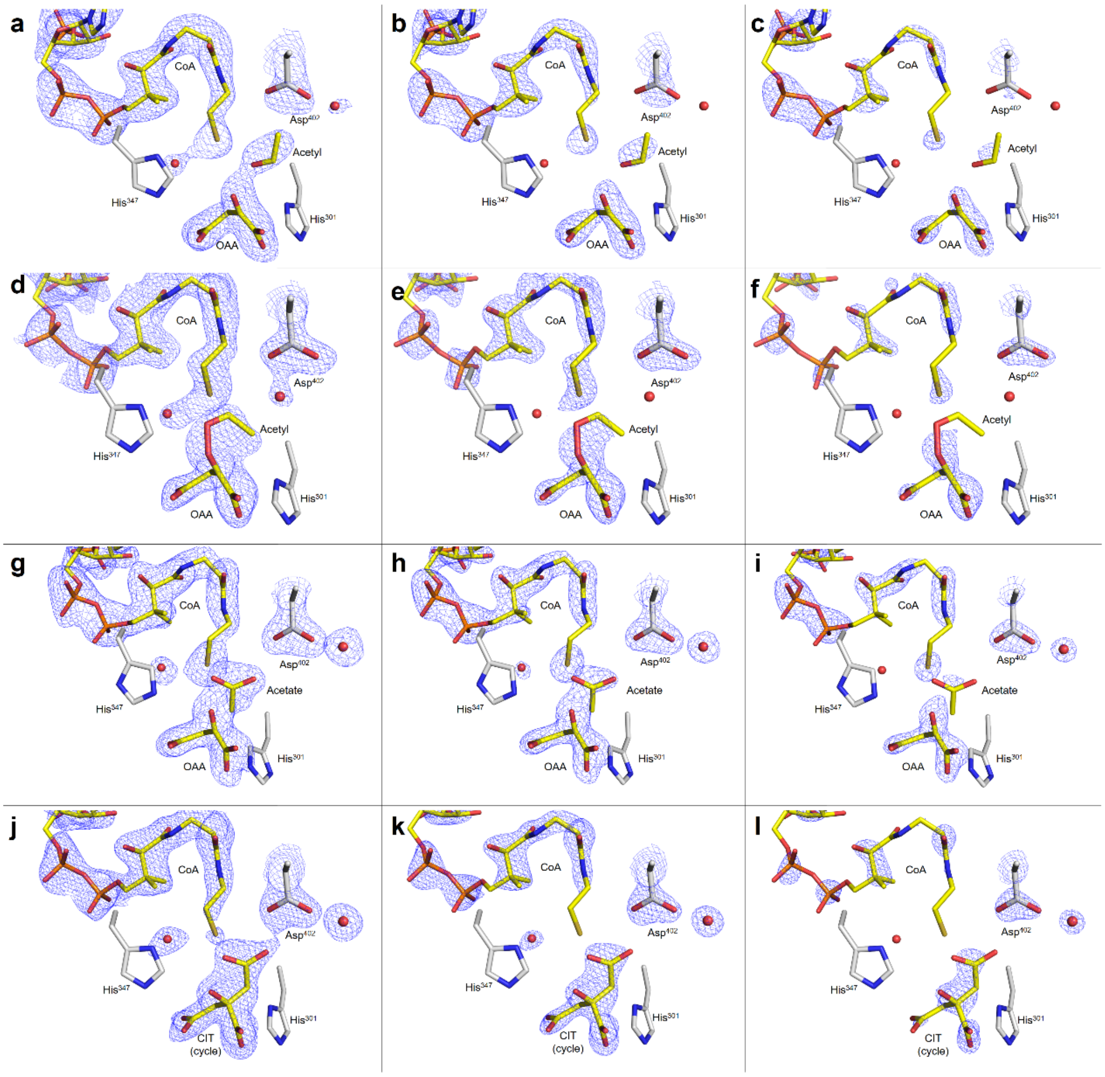


**Supplementary Fig. 16 | Structures of intermediates in active site of *h*CS.**

**a,** Acetyl analogue as intermediate with oxaloacetate (OAA) and CoA (yellow) in *h*CS active site (gray) (PDB: 8ZVM). 2Fo-Fc omit map are shown as blue mesh in 1.0σ. **b,** Same to **a** with 1.5σ map. **c,** Same to **a** with 2.0σ map. **d,** Acetyl analogue forming peroxide bond with oxaloacetate in *h*CS active site (PDB: 8ZVR) with 1.0σ map. **e,** Same to **d** with 1.5σ map. **f,** Same to **d** with 2.0σ map. **g,** Acetate analogue in *h*CS active site (PDB: 8ZVT) with 1.0σ map. **h,** Same to **g** with 1.5σ map. **i,** Same to **g** with 2.0σ map. **j,** Citrate forming five-member cycle in *h*CS active site (PDB: 8ZVU) with 1.0σ map. **k,** Same to **j** with 1.5σ map. **l**, Same to **j** with 2.0σ map.

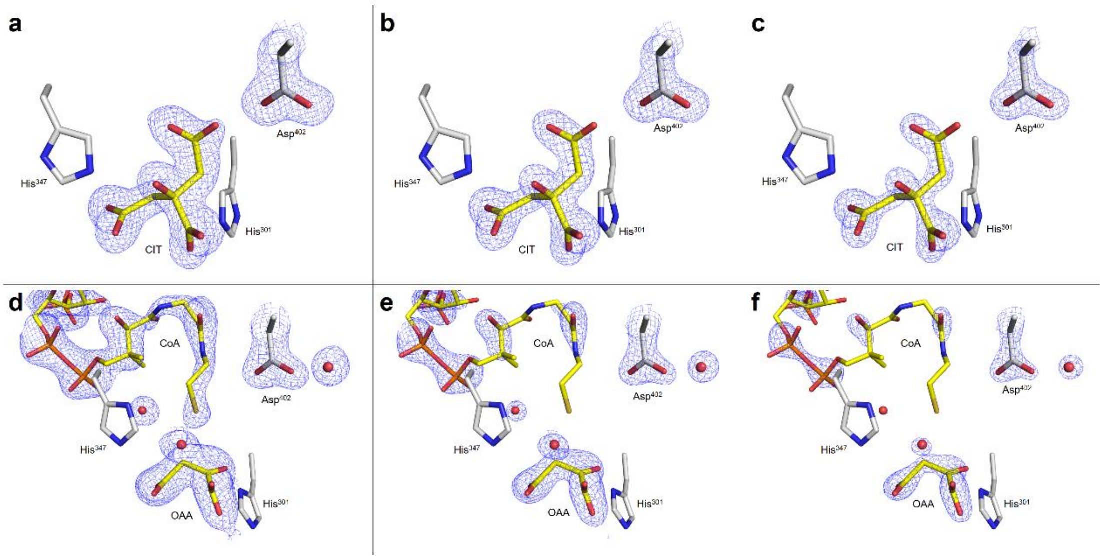


**Supplementary Fig. 17 | Structures of products in active site of *h*CS.**

**a,** Citrate (CIT, yellow) as product binding in *h*CS active site (gray) (PDB: 8ZVV). 2Fo-Fc omit map are shown as blue mesh in 1.0σ. **b,** Same to **a** with 1.5σ map. **c,** Same to **a** with 2.0σ map. **d,** Oxaloacetate with an hydroxyl group and CoA in *h*CS active site (PDB: 8ZVW) with 1.0σ map. **e,** Same to **d** with 1.5σ map. **f,** Same to **d** with 2.0σ map.

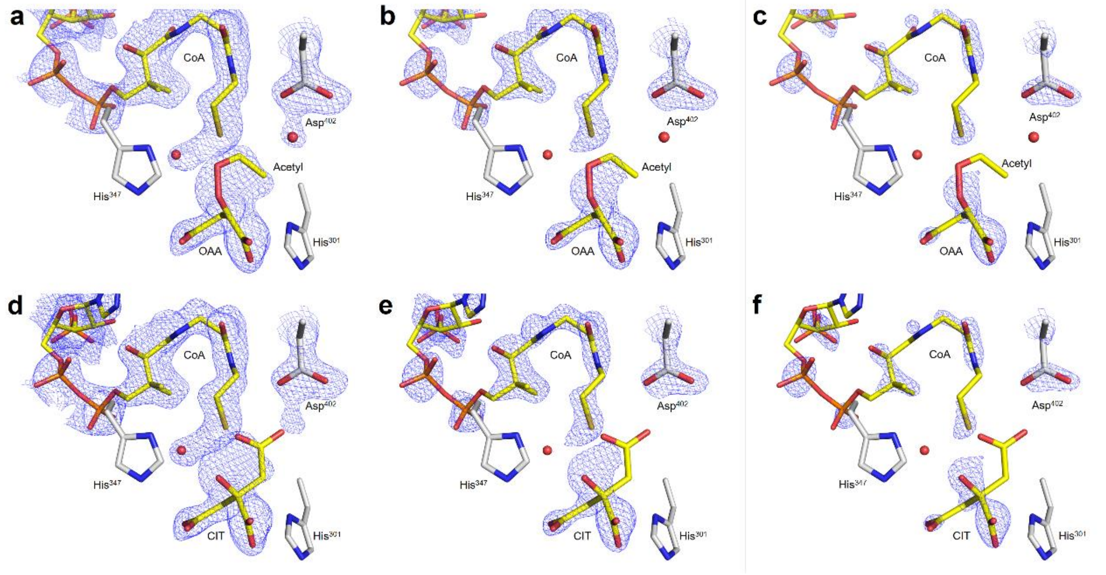


**Supplementary Fig. 18 | The electron density more suitable for oxaloacetate and acetyl group rather than citrate in *h*CS-intermediate 2 structure.**

**a,** Acetyl analogue as intermediate with oxaloacetate (OAA) and CoA (yellow) in *h*CS active site (gray) (PDB: 8ZVR). 2Fo-Fc omit map are shown as blue mesh in 1.0σ. **b,** Same to **a** with 1.5σ map. **c,** Same to **a** with 2.0σ map. **d,** Same 2Fo-Fc omit map to **a** in 1.0σ, with citrate as ligand. **e,** Same to **d** with 1.5σ map. **f,** Same to **d** with 2.0σ map.

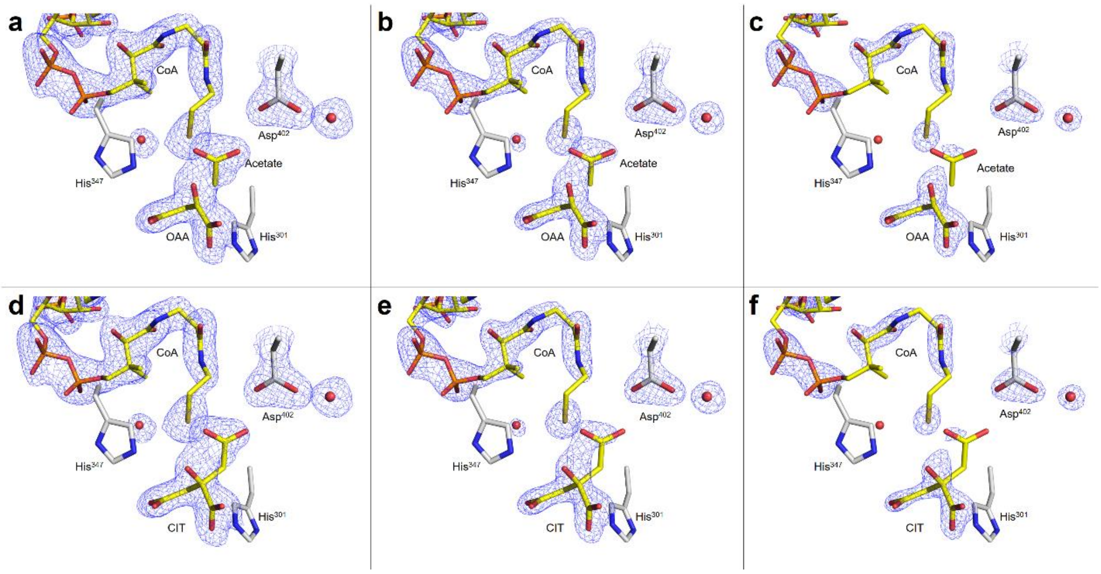


**Supplementary Fig. 19 | The electron density more suitable for oxaloacetate and acetate analogue rather than citrate in *h*CS-intermediate 3 structure.**

**a,** Acetate analogue as intermediate with oxaloacetate (OAA) and CoA (yellow) in *h*CS active site (gray) (PDB: 8ZVT). 2Fo-Fc omit map are shown as blue mesh in 1.0σ. **b,** Same to **a** with 1.5σ map. **c,** Same to **a** with 2.0σ map. **d,** Same 2Fo-Fc omit map to **a** in 1.0σ, with citrate as ligand. **e,** Same to **d** with 1.5σ map. **f,** Same to **d** with 2.0σ map.

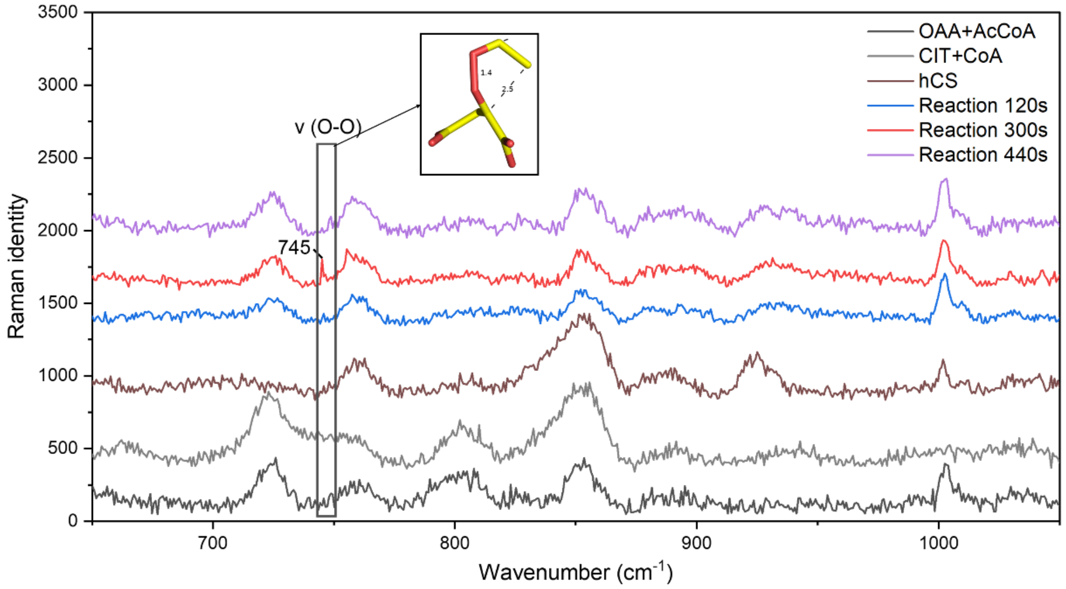


**Supplementary Fig. 20 | Raman spectra of *h*CS crystal reacting different times showing 745 cm$^{-1}$ band.**

Raman spectra of *h*CS crystal after mixed with substrates for 120s (blue), 300s (red) and 440s (purple), range of 650-1050 cm$^{-1}$. The concerned band appears at 745 cm$^{-1}$ in spectrum of 300s and labeled. The corresponding intermediate structure are shown in yellow sticks. Raman spectra of 30 mM oxaloacetate and 50 mM acetyl-CoA (black), 30 mM citrate and 50 mM CoA (gray) and *h*CS (brown) is shown as control.

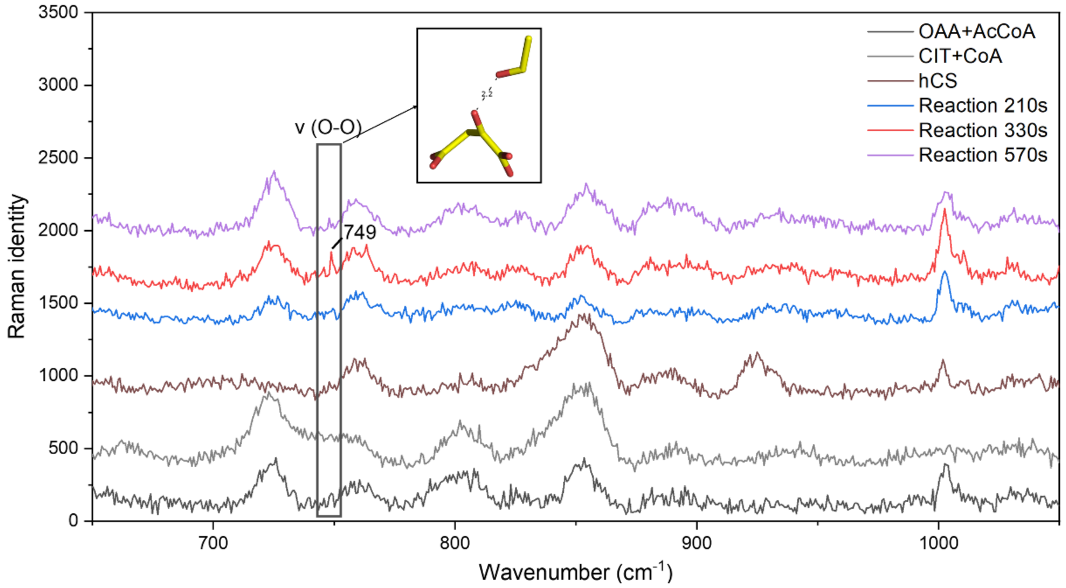


**Supplementary Fig. 21 | Raman spectra of *h*CS crystal reacting different times showing 749 cm$^{-1}$ band.**

Raman spectra of *h*CS crystal (different from that in **Supplementary Fig. 20**) after mixed with substrates for 210s (blue), 330s (red) and 570s (purple), range of 650-1050 cm$^{-1}$. The concerned band appears at 749 cm$^{-1}$ in spectrum of 330s and labeled. The corresponding intermediate structure are shown in yellow sticks. Raman spectra of 30 mM oxaloacetate and 50 mM acetyl-CoA (black), 30 mM citrate and 50 mM CoA (gray) and *h*CS (brown) is shown as control.

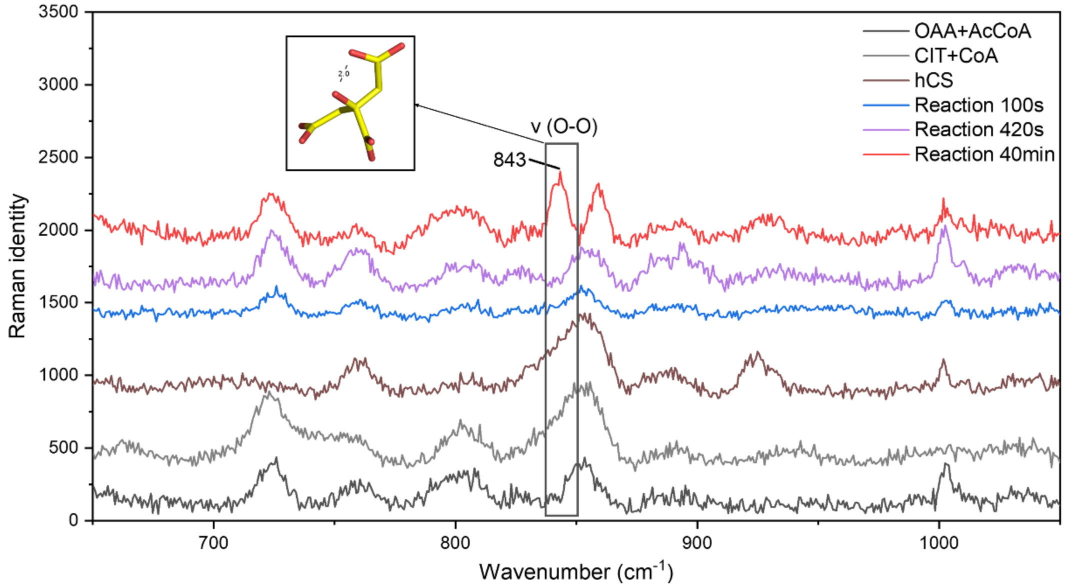


**Supplementary Fig. 22 | Raman spectra of *h*CS crystal reacting different times showing 843 cm$^{-1}$ band.**

Raman spectra of *h*CS crystal (different from that in **Supplementary Fig. 20 and 21**) after mixed with substrates for 100s (blue), 420s (purple) and 40 min (red), range of 650-1050 cm$^{-1}$. The concerned band appears at 843 cm$^{-1}$ in spectrum of 40min and labeled. The corresponding intermediate structure are shown in yellow sticks. Raman spectra of 30 mM oxaloacetate and 50 mM acetyl-CoA (black), 30 mM citrate and 50 mM CoA (gray) and *h*CS (brown) is shown as control.

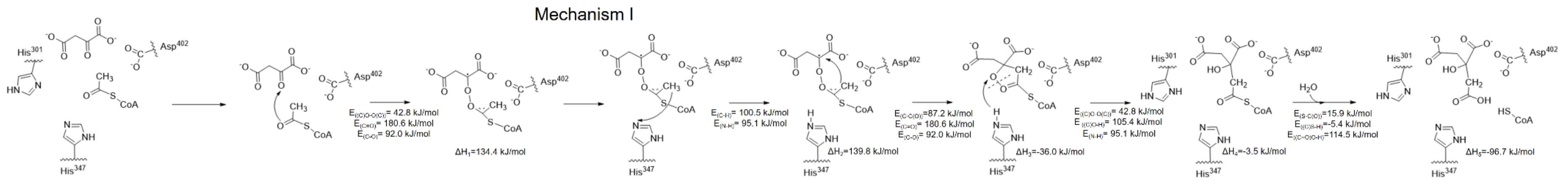


**Supplementary Fig. 23 | Energy change in catalytic mechanism I.** $\Delta H=E_{(bond\text{-}formation)}-E_{(bond\text{-}broken)}$. $\Delta H_1=(E_{(C=O)}$[26]$-E_{(C\text{-}O)}$[27]$)\times2-E_{((C)O\text{-}O(C))}$[28]. $\Delta H_2=\Delta H_1+E_{((C)C\text{-}H)}$[29]$-E_{((IMD)N\text{-}H)}$[30]. $\Delta H_3=\Delta H_2-E_{(C\text{-}C(O))}$[26]$-(E_{(C=O)}$[26]$-E_{(C\text{-}O)}$[27]$)$. $\Delta H_4=\Delta H_3+E_{((C)O\text{-}O(C))}$[28]$+E_{((IMD)N\text{-}H)}$[30]$-E_{((C)O\text{-}H)}$[31]. $\Delta H_5=\Delta H_4+E_{(S\text{-}C(O))}$[32-33]$-E_{((C)S\text{-}H)}$[32-33]$-E_{((C=O)O\text{-}H)}$[34].

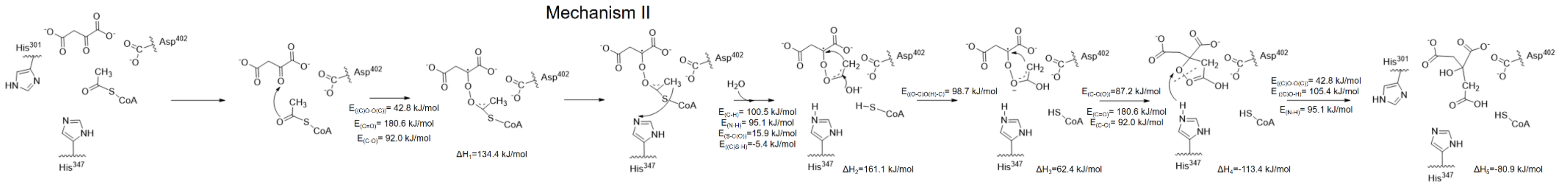


**Supplementary Fig. 24 | Energy change in catalytic mechanism II.** $\Delta H_1=(E_{(C=O)}$[26]$-E_{(C\text{-}O)}$[27]$)\times2-E_{((C)O\text{-}O(C))}$[28]. $\Delta H_2=\Delta H_1+E_{((C)C\text{-}H)}$[29]$-E_{((IMD)N\text{-}H)}$[30]$+E_{(S\text{-}C(O))}$[32-33]$-E_{((C)S\text{-}H)}$[32-33]. $\Delta H_3=\Delta H_2-E_{((O\text{-}C)C\text{-}O(H))}$[35]. $\Delta H_4=\Delta H_3-E_{(C\text{-}C(O))}$[26]$-(E_{(C=O)}$[26]$-E_{(C\text{-}O)}$[27]$)$. $\Delta H_5=\Delta H_4+E_{((C)O\text{-}O(C))}$[28]$+E_{((IMD)N\text{-}H)}$[30]$-E_{((C)O\text{-}H)}$[31].

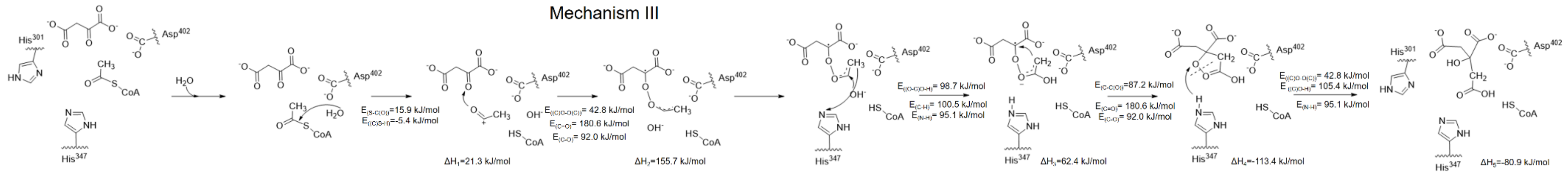


**Supplementary Fig. 25 | Energy change in catalytic mechanism III.**

$\Delta H_1=E_{(S\text{-}C(O))}$[32-33]$-E_{((C)S\text{-}H)}$[32-33]. $\Delta H_2=\Delta H_1+(E_{(C=O)}$[26]$-E_{(C\text{-}O)}$[27]$)\times 2-E_{((C)O\text{-}O(C))}$[28]. $\Delta H_3=\Delta H_2+E_{((C)C\text{-}H)}$[29]$-E_{((IMD)N\text{-}H)}$[30]$-E_{((O\text{-}C)C\text{-}O(H))}$[35]. $\Delta H_4=\Delta H_3-E_{(C\text{-}C(O))}$[26]$-(E_{(C=O)}$[26]$-E_{(C\text{-}O)}$[27]$)$. $\Delta H_5=\Delta H_4+E_{((C)O\text{-}O(C))}$[28]$+E_{((IMD)N\text{-}H)}$[30]$-E_{((C)O\text{-}H)}$[31].

**Table S1. Assigned signals of DMPO-radical adduct in EPR spectra.**

| Sample component | radical species | Peak No.[a] | g-factor [b] | hyperfine splitting values: $A_N$ (G) [c] | hyperfine splitting values: $A_H^{\beta}$ (G) [d] | Intensity ratio [e] |
|---|---|---|---|---|---|---|
| *Da*CS+CIT+ CoA | alkyl radical | 6-line (1, 3, 5, 6, 8, 10) | 2.006 | 15.58 (15~17)[f] | 22.41 (21~23)[f] | 1.27:1.35:1.00:1.00:1.09:1.09 (1:1:1:1:1:1)[f] |
| | hydroxyl radical | 4-line (2, 4, 7, 9) | 2.006 | 14.84 (14~15)[f] | 14.81 (14~15)[f] | 1.00:1.92:1.90:1.14 (1:2:2:1)[f] |
| CIT+CoA | alkyl radical | 6-line (1, 3, 5, 6, 8, 10) | 2.006 | 15.62 | 22.40 | 1.16:1.18:1.07:1.08:1.06:1.00 |
| | hydroxyl radical | 4-line (2, 4, 7, 9) | 2.006 | 14.86 | 14.85 | 1.29:2.02:1.80:1.00 |
| CIT | alkyl radical | 6-line (1, 3, 5, 6, 8, 10) | 2.006 | 15.49 | 22.80 | 1.20:1.33:1.24:1.00:1.41:1.09 |
| | hydroxyl radical | 4-line (2, 4, 7, 9) | 2.006 | 14.84 | 14.81 | 1.00:1.82:1.79:1.01 |
| CoA | alkyl radical | 6-line (1, 3, 5, 6, 8, 10) | 2.006 | 15.63 | 22.66 | 1.06:1.19:1.28:1.05:1.16:1.00 |
| | hydroxyl radical | 4-line (2, 4, 7, 9) | 2.006 | 14.84 | 14.81 | 1.00:1.78:1.63:1.00 |
| | Possible CoA thiyl radical | 6-line (2, 3, 5, 6, 8, 9) | 2.006 | 13.30 | 20.33 | 1.44:1.13:1.21:1.00:1.10:1.43 |
| | | 6-line (2, 4, 5, 6, 7, 9) | 2.006 | 13.30 | 16.38 | 1.44:2.55:1.21:1.00:2.34:1.43 |
| | | 6-line (2, 3, 4, 7, 8, 9) | 2.006 | 14.84 | 18.79 | 1.30:1.02:2.31:2.12:1.00:1.30 |
| | | 6-line (1, 4, 5, 6, 7, 10) | 2.006 | 15.63 | 18.70 | 1.06:2.69:1.28:1.05:2.47:1.00 |
| | | 6-line (1, 3, 4, 7, 8, 10) | 2.006 | 17.17 | 21.12 | 1.06:1.19:2.69:2.47:1.16:1.00 |
| CoA+$H_2O_2$+ Peroxidase[36] | CoA-thiyl radical | 4-line | - | 15.4 | 16.2 | 1:1:1:1 |
| Cys+$H_2O_2$+ Peroxidase[37] | Cysteine-thiyl radical | 6-line | - | 15.3 | 17.0 | 1:1:1:1:1:1 |

**a,** In each EPR spectrum of *Da*CS reverse reaction system or substrate, ten peaks were picked out and assigned as signal of DMPO-radical adduct. These peaks were numbered from smaller magnetic field to large one.

**b,** The spectroscopic splitting factor (g factor) $g = h\nu/\beta B$, in which Planck constant $h=6.626068\times10^{-34}$ $m^2\cdot kg/s$, Bohr magneton $\beta=9.27\times10^{-24}$ $A\cdot m^2$, the frequency of microwave $\nu=9.438$ GHz in this experimental conditions, and the magnetic field B is the mean value of the horizontal coordinate for the characteristic peak's intersections with the baseline.

**c,** For alkyl radical, hyperfine splitting value $A_N$ is the distance between the 1$^{st}$ and the 2$^{nd}$ line, or the 2$^{nd}$ and the 4$^{th}$ line, or the 3$^{rd}$ and the 5$^{th}$ line, or the 5$^{th}$ and the 6$^{th}$ line. For hydroxyl radical, $A_N = A_H^{\beta}$, is distance between the mean of the 3$^{rd}$ and the 4$^{th}$ line and the mean of the 1$^{st}$ and the 2$^{nd}$ line.

**d,** For alkyl radical, hyperfine splitting value $A_H^{\beta}$ is the distance between the 2$^{nd}$ and the 5$^{th}$ line. For hydroxyl radical, $A_H^{\beta} = A_N$, is the distance between the 2$^{nd}$ and the 3$^{rd}$ line.

**e,** Intensity ratio was the ratio of peak heights for peaks assigned as DMPO-radical adduct signal.

**f**, Values in parentheses are the theoretical value of hyperfine splitting constant and intensity ratio for DMPO-radical adduct.

### Table S2. Data collection and refinement statistics of *Da*CS and *Tt*CS.

| Datasets | *Da*CS-citrate | DaCS-intermediate | *Da*CS-oxaloacetate | TtCS-citrate/CoA | *Tt*CS-intermediate | *Tt*CS-oxaloacetate/ acetyl-CoA |
|---|---|---|---|---|---|---|
| PDB ID | 8XU2 | 8XXC | 8XU1 | 8XXD | 8XXE | 8XXF |
| **Data collection** | | | | | | |
| Space group | *H*3 | *H*3 | *C*2 | $P4_32_12$ | $P4_32_12$ | $P2_12_12_1$ |
| Cell dimensions | | | | | | |
| *a*, *b*, *c* (Å) | 362.33, 362.33, 77.89 | 361.42, 361.42, 77.74 | 155.10, 111.28, 99.56 | 109.56, 109.56, 265.20 | 109.91, 109.91, 265.09 | 108.49, 131.76, 207.24 |
| a, b, γ (°) | 90.00, 90.00, 120.00 | 90.00, 90.00, 120.00 | 90.00, 116.07, 90.00 | 90.00, 90.00, 90.00 | 90.00, 90.00, 90.00 | 90.00, 90.00, 90.00 |
| Resolution (Å) | 50.00-2.18 (2.22-2.18) | 50.00-2.03 (2.07-2.03) | 45.11-2.46 (2.52-2.46) | 50.63-2.32 (2.38-2.32) | 34.46-2.08 (2.13-2.08) | 30.15-2.33 (2.39-2.33) |
| $R_{merge}$ | 0.129 (0.567) | 0.185 (0.983) | 0.182 (1.366) | 0.182 (1.366) | 0.181 (1.313) | 0.319 (2.213) |
| $R_{meas}$ | 0.137 (0.606) | 0.199 (1.060) | 0.255 (1.907) | 0.190 (1.419) | 0.189 (1.363) | 0.349 (2.606) |
| $CC_{1/2}$ | 0.991 (0.869) | 0.989 (0.680) | 0.981 (0.311) | 0.998 (0.854) | 0.999 (0.896) | 0.983 (0.449) |
| $I / \sigma I$ | 32.0 (4.7) | 14.5 (2.0) | 6.7 (1.3) | 14.9 (3.4) | 19.3 (4.8) | 5.9 (0.9) |
| Data completeness (%) | 97.5 (96.4) | 99.1 (99.8) | 99.1 (99.1) | 99.9 (100.0) | 100.0 (100.0) | 99.9 (99.9) |
| Redundancy | 8.2 | 7.3 | 3.3 | 25.0 | 25.7 | 11.7 |
| **Refinement** | | | | | | |
| No. reflections | 181,722 | 223,879 | 51,987 | 67,128 | 93,238 | 120,779 |
| $R_{work}$ / $R_{free}$ | 0.176 / 0.203 | 0.198 / 0.239 | 0.213 / 0.258 | 0.190 / 0.243 | 0.173 / 0.217 | 0.189 / 0.245 |
| R.m.s. deviations | | | | | | |
| Bond lengths (Å) | 0.007 | 0.007 | 0.008 | 0.008 | 0.008 | 0.007 |
| Bond angles (°) | 1.314 | 1.428 | 1.470 | 1.471 | 1.555 | 1.712 |
| No. atoms | 29117 | 29531 | 10424 | 11305 | 11484 | 22328 |
| Protein | 27491 | 27401 | 10275 | 10725 | 10712 | 21250 |
| Citrate | 104 | 78 | - | 13 | - | - |
| Oxaloacetate | - | 18 | 27 | 18 | 27 | 45 |
| Acetate analogue | - | 8 | - | - | - | - |
| Acetyl group | - | - | - | - | 3 | - |
| Coenzyme A | - | - | - | 96 | 144 | 48 |
| Acetyl-coenzyme A | - | - | - | - | - | 102 |
| Water | 1522 | 2026 | 122 | 183 | 598 | 883 |
| *B*-factors | 39.5 | 35.8 | 59.5 | 53.6 | 41.6 | 39.3 |
| Protein | 39.8 | 35.6 | 59.7 | 53.4 | 41.0 | 39.4 |
| Citrate | 35.4 | 27.8 | - | 69.4 | - | - |
| Oxaloacetate | - | 34.4 | 64.3 | 77.5 | 59.5 | 56.4 |
| Acetate analogue | - | 49.9 | - | - | - | - |
| Acetyl group | - | - | - | - | 40.4 | - |
| Coenzyme A | - | - | - | 79.6 | 70.8 | 48.2 |
| Acetyl-coenzyme A | - | - | - | - | - | 63.6 |
| Water | 34.8 | 40.0 | 40.5 | 47.4 | 44.5 | 32.7 |

Values in parentheses are for the highest resolution shell.

**Table S3. Sequence and structure similarity between *Da*CS and other five citrate synthases**

| Protein | Species | Residues (without signal peptide) | Sequence identity % | PDB code | Ligand | C-α r.m.s.d (Å). |
|---|---|---|---|---|---|---|
| *Da*CS | *Desulfurella acetivorans* | 1-436 | 100.00 | 8XU2 | Citrate | - |
| *Tt*CS | *Thermosulfidibacter takaii* | 1-442 | 59.26 | 8XXD | Citrate and coenzyme A | 1.1 |
| *h*CS | *Homo sapiens (Human)* | 28-466 | 50.58 | 5UZR | - | 1.4 |
| *At*CS | *Arabidopsis thaliana* | 17-474 | 53.12 | 6K5V | - | 1.3 |
| *Tp*CS | *Thermoplasma acidophilum* | 1-385 | 23.20 | 4YBO | - | 2.3 |
| *Sc*CS | *Saccharolobus solfataricus* | 1-377 | 26.76 | 1O7X | - | 2.3 |

Sequence alignment (without signal peptide of *h*CS and *At*CS) and percent identity matrix were calculated using Clustal Omega (v2.1)[19-21] on EBI web server[22] (www.ebi.ac.uk/Tools/msa/clustalo). Structure similarity is represented by root mean square deviation (R.M.S.D) which were generated by DALI[38].

**Table S4. Sequence and structure similarity between *Tt*CS and other five citrate synthases.**

| Protein | Species | Residues (without signal peptide) | Sequence identity % | PDB code | Ligand | C-α r.m.s.d (Å). |
|---|---|---|---|---|---|---|
| *Tt*CS | *Thermosulfidibacter takaii* | 1-442 | 100.00 | 8XXD | Citrate and coenzyme A | - |
| *Da*CS | *Desulfurella acetivorans* | 1-436 | 59.26 | 8XU2 | Citrate | 1.1 |
| *h*CS | *Homo sapiens (Human)* | 28-466 | 53.56 | 5UZR | - | 1.3 |
| *At*CS | *Arabidopsis thaliana* | 17-474 | 54.59 | 6K5V | - | 1.1 |
| *Tp*CS | *Thermoplasma acidophilum* | 385 | 21.49 | 4YBO | - | 2.3 |
| *Sc*CS | *Saccharolobus solfataricus* | 1-377 | 24.80 | 1O7X | - | 2.3 |

Sequence alignment (without signal peptide of *h*CS and *At*CS) and percent identity matrix were calculated using Clustal Omega (v2.1)[19-21] on EBI web server[22] (www.ebi.ac.uk/Tools/msa/clustalo). Structure similarity is represented by root mean square deviation (R.M.S.D) which were generated by DALI[38].

**Table S5. Maximum forward velocity for *Da*CS and *Tt*CS under aerobic or anaerobic conditions**

| Enzyme | Conditions | | $V_{max}$ (μmol/min/mg) | |
|---|---|---|---|---|
| | Gas environment | Temperature (°C) | Oxaloacetate | Acetyl-CoA |
| *Da*CS | air | 50 | 46.00±0.76 | |
| *Da*CS[1] | $N_2$[1] | 55[1] | 131±3[1] | 157±4[1] |
| *Tt*CS | air | 70 | 58.92±11.12 | |
| *Tt*CS[17] | $N_2$[17] | 70[17] | 239±12[17] | 298±13[17] |

Referenced Data in $N_2$ are mean ± s.d. of three technical replicates, while data in air are mean ± s.e. of three technical replicate.

### Table S6. Data collection and refinement statistics of *h*CS.

| Datasets | *hCS*-oxaloacetate | *h*CS-oxaloacetate/ acetyl-CoA | *h*CS-intermediate 1 | *h*CS-intermediate 2 | *h*CS-intermediate 3 | *h*CS-intermediate 4 | *h*CS-citrate | *h*CS-byproduct |
|---|---|---|---|---|---|---|---|---|
| PDB ID | 8ZVL | 8ZW1 | 8ZVM | 8ZVR | 8ZVT | 8ZVU | 8ZVV | 8ZVW |
| **Data collection** | | | | | | | | |
| Space group | *P*1 | *P*1 | *P*1 | *P*1 | $P4_32_12$ | *P*1 | $P\ 12_11$ | $P4_32_12$ |
| Cell dimensions | | | | | | | | |
| *a*, *b*, *c* (Å) | 57.60, 78.01, 103.12 | 57.52, 59.79, 72.58 | 58.21, 59.82, 73.24 | 57.92, 59.73, 73.28 | 57.84, 57.84, 250.36 | 57.91, 59.69, 73.43 | 57.55, 194.33, 69.37 | 57.37, 57.37, 250.67 |
| a, b, γ (°) | 70.75, 81.37, 74.74 | 102.52, 98.23, 118.76 | 101.72, 98.66, 116.55 | 101.58, 98.87, 116.40 | 90.00, 90.00, 90.00 | 101.45, 98.91, 116.47 | 90.00, 90.40, 90.00 | 90.00, 90.00, 90.00 |
| Resolution (Å) | 32.68-2.05 (2.10-2.05) | 67.83-2.03 (2.14-2.03) | 68.94-1.97 (2.17-1.97) | 68.97-1.80 (1.90-1.80) | 31.68-2.05 (2.10-2.05) | 50.93-1.78 (1.88-1.78) | 97.17-1.59 (1.68-1.59) | 29.10-1.82 (1.87-1.82) |
| $R_{\text{merge}}$ | 0.151 (0.828) | 0.114 (0.841) | 0.115 (0.603) | 0.140 (1.163) | 0.262 (1.586) | 0.104 (1.109) | 0.134 (1.272) | 0.227 (2.068) |
| $R_{meas}$ | 0.181 (1.012) | 0.151 (1.134) | 0.149 (0.793) | 0.180 (1.508) | 0.268 (1.617) | 0.128 (1.345) | 0.150 (1.456) | 0.232 (2.218) |
| $CC_{1/2}$ | 0.990 (0.423) | 0.978 (0.223) | 0.981 (0.309) | 0.895 (0.501) | 0.998 (0.754) | 0.968 (0.404) | 0.987 (0.323) | 0.995 (0.348) |
| $I / \sigma I$ | 5.7 (1.8) | 1.6 (0.8) | 2.9 (1.5) | 5.3 (1.3) | 12.6 (2.7) | 6.8 (2.0) | 6.5 (1.8) | 8.8 (1.2) |
| Data completeness (%) | 96.4 (95.9) | 94.2 (96.0) | 84.6 (44.5) | 88.3 (83.3) | 99.9 (100.0) | 90.3 (89.6) | 99.1 (97.7) | 100.0 (100.0) |
| Redundancy | 3.4 | 2.0 | 2.0 | 2.3 | 24.7 | 2.8 | 4.8 | 22.4 |
| **Refinement** | | | | | | | | |
| No. reflections | 93672 | 41416 | 34230 | 61068 | 26468 | 66996 | 191015 | 36998 |
| $R_{\text{work}} / R_{\text{free}}$ | 0.197 / 0.258 | 0.236 / 0.299 | 0.221 / 0.268 | 0.223 / 0.274 | 0.204 / 0.269 | 0.186 / 0.233 | 0.244 / 0.295 | 0.226 / 0.298 |
| R.m.s. deviations | | | | | | | | |
| Bond lengths (Å) | 0.008 | 0.003 | 0.003 | 0.008 | 0.007 | 0.009 | 0.006 | 0.007 |
| Bond angles (°) | 1777 | 1.118 | 1.173 | 1.727 | 1.630 | 1.756 | 1.413 | 1.698 |
| No. atoms | 14280 | 7082 | 6998 | 7257 | 3637 | 7252 | 14604 | 3569 |
| Protein | 13698 | 6819 | 6816 | 6827 | 3424 | 6837 | 13688 | 3416 |

| | | | | | | | | |
|---|---|---|---|---|---|---|---|---|
| Citrate | - | 13 | - | 13 | - | 13 | 52 | - |
| Oxaloacetate | 27 | 9 | 9 | 9 | 9 | 9 | - | 9 |
| Acetyl analogue | - | - | 3 | 3 | - | - | - | - |
| Acetate analogue | - | - | - | - | 4 | - | - | - |
| Coenzyme A | - | 48 | 48 | 48 | 48 | 48 | - | 48 |
| Acetyl-CoA | - | 51 | - | - | - | - | - | - |
| Water | 555 | 142 | 122 | 357 | 152 | 345 | 864 | 96 |
| *B*-factors | 42.3 | 50.5 | 45.3 | 40.6 | 34.9 | 37.7 | 31.3 | 46.5 |
| Protein | 42.2 | 50.2 | 45.2 | 40.3 | 34.5 | 37.2 | 30.9 | 46.0 |
| Citrate | - | 44.1 | - | - | - | 48.2 | 28.9 | - |
| Oxaloacetate | 47.1 | 63.3 | 39.4 | 37.3 | 36.1 | 57.3 | - | 52.9 |
| Acetyl analogue | - | - | 60.1 | 54.5 | - | - | - | - |
| Acetate analogue | - | - | - | - | 60.8 | - | - | - |
| Coenzyme A | - | 62.2 | 68.5 | 74.8 | 53.2 | 80.8 | - | 85.3 |
| Acetyl-CoA | - | 93.3 | - | - | - | - | - | - |
| Water | 44.8 | 44.7 | 39.0 | 41.1 | 35.7 | 39.7 | 37.9 | 43.0 |

Values in parentheses are for the highest resolution shell.
